# Adenocarcinoma cell mechanobiology is altered by the loss modulus of the surrounding extracellular matrix

**DOI:** 10.64898/2026.02.04.703912

**Authors:** Ariell M. Smith, Brandon M. Pardi, Isaac Sousa, Arvind Gopinath, Roberto Andresen Eguiluz

## Abstract

Elastic and viscoelastic properties of extracellular matrices (ECM) are known to regulate cellular behavior and mechanosensation differently, with implications for morphogenesis, wound healing, and pathophysiology. Most *in vitro* cellular processes, including cell migration, are studied on linear-elastic substrates to mimic extracellular matrices. However, most tissues are viscoelastic and display a loss modulus (*G*⍰⍰) that may be 10-20% of their storage modulus (*G*⍰) under biophysically relevant conditions. Recent research has shown that cells can distinguish between elastic and viscoelastic ECM, leading to alterations in their cellular morphology, migration rates, and contractility. Here, we present a protocol for creating PAH-based model ECMs that enables the fabrication of viscoelastic substrates with storage moduli similar to those of their elastic counterparts. To explore how *G*⍰⍰ influences epithelial cell mechanobiology, we fabricated tunable viscoelastic model ECMs with *G*⍰ of 3 kPa, 8 kPa, and 12 kPa, and for each, independently tuned *G*⍰⍰ values to approximately 300 Pa, 500 Pa, and 700 Pa, respectively. We found that A549 cells cultured on stiff elastic model ECMs migrated ∼30% slower and formed larger focal adhesions compared to their viscoelastic counterparts. Conversely, A549 cells on intermediate viscoelastic model ECMs exhibited a ∼54% reduction in migration speed, with no significant difference in focal adhesion size relative to their elastic counterparts. These findings highlight the complex interplay between substrate (ECM) elastic and viscoelastic properties in regulating epithelial cell mechanobiology and emphasize the importance of time-dependent matrix mechanics in governing epithelial responses.

## 1. Introduction

The extracellular matrix (ECM) is a macromolecular scaffold that provides mechanical support and structure to cells.^1–4^ The mechanical properties of the ECM depend on the concentration and ratio of proteins,^1,2^ fiber orientation,^4^ molecular conformation,^3^ among other factors.^5^ This is evidenced during aging or pathophysiology, conditions in which ECM protein ratios, orientation, and confirmation of the molecular constituents change.^3,5,6^ These changes lead to significant alterations in the mechanical properties of the ECM,^7^ including elastic and viscoelastic properties, such as stiffness or relaxation times.^8^ Decades of research have established that elasticity alone can regulate cellular behavior. For example, adherent cells, such as fibroblasts (3T3-Swiss) and epithelial (normal rat kidney) cells, on rigid model EMCs exhibit larger, longer, and more mature focal adhesions than on compliant ECMs, leading to larger cytoskeletal size, increased cell spreading area, and increased cell proliferation.^9–11^ Substrate stiffness also mediates cell differentiation. For example, mesenchymal stem cells on softer ECMs (stiffness 0.1-1 kPa, corresponding to, for instance, brain-like tissue) become neurogenic, while those on rigid ECMs (stiffness 25-40 kPa, seen in bone-like tissue) turn osteogenic.^12,13^ Yet another example is the organization of the human umbilical vascular endothelial (HUVECs) network. In this instance, cells communicate mechanically by applying strain fields to the substrates, facilitating and coordinating the formation of a network. This mechanical signaling has been demonstrated for cells on substrates that are neither too stiff nor too compliant.^9,14,15^ Therefore, it is clear that if the mechanical environment is altered, the cues sensed by the cells modify mechanotransductive signaling pathways. This is the case for the mechanosensitive signaling pathway YAP/TAZ, which activates integrins, focal adhesions, the cytoskeleton, and transmits signals to the Hippo pathway, thereby altering cell proliferation, growth,^16^ and fibrosis.^17,18^

Several studies in mechanobiology have focused mainly on the elastic properties of the ECM; however, ECMs are intrinsically viscoelastic.^19,20^ Viscoelastic materials exhibit a complex response to stress or strain, encompassing instantaneous as well as time-dependent responses.^21,22^ Temporal responses are commonly characterized by relaxation time constants, the loss modulus (encapsulating the non-elastic, dissipative time-dependent responses), and the storage modulus (quantifying elastic, instantaneous responses). Native tissue elastic moduli range from extremely soft (∼0.5 kPa, fat tissue)^23^ to very rigid (1-5 GPa, bone tissue);^24^ loss moduli meanwhile, depending on tissue type, can vary from 10-20% of the associated elastic moduli under biophysically relevant conditions.^21^ It has been shown that model ECMs with similar elastic moduli but distinct (different) loss moduli affect cells in a cell-line-specific manner.^25,26^ These include differences in cell spreading,^27–29^ cell migration,^21,30^ polarization, and differentiation,^21,31,32^ among others. Typically, alginate or alginate-based gels have been used to fabricate a broad range of tunable elastic and viscoelastic model ECMs to investigate cell mechanobiology.^29,33–36^ 3T3 mouse fibroblast cells seeded on alginate model ECMs with Young’s modulus of 1.4 kPa and varying loss moduli, exhibited larger cell-spreading areas and stress fiber formation on viscoelastic model ECMs than on linear elastic model ECMs.^29^ Conversely, human mesenchymal stem cells seeded on alginate-PEG hydrogels with a Young’s modulus of approximately 3 kPa and varying viscoelasticity showed a larger spread area on elastic (faster stress relaxation) compared to viscoelastic (slower stress relaxation) model ECMs.^28^ Cell-specific responses are not unique to cells on alginate substrates. Differences in cellular responses have been illustrated using elastic and viscoelastic polyacrylamide-based (PAH-based) model ECMs. For example, Huh7 and primary human hepatocytes were explored on elastic (*G*□ = 5 kPa, *G*□□ = 0 Pa) and viscoelastic (*G*□ = 5 kPa, *G*□□ = 600 Pa) polyacrylamide hydrogel (PAH) substrates and exhibited opposite mechanobiology responses. Huh7 displayed increased cell area, speed, and longer protrusion lengths on viscoelastic model ECMs. Conversely, hepatocytes displayed decreased cell area and speed on viscoelastic model ECMs.^37^

PAHs are another commonly used platform for cell mechanobiology studies, as model ECMs can be readily fabricated across a wide range of physiologically relevant elasticities. To introduce dissipative effects and increase the loss moduli of these elastic model ECMs, linear acrylamide chains can be embedded into the elastic PAH network.^21,32^ Using a PAH platform of tunable viscoelasticity, it is reported that human mammary epithelial cells (MCF10A) seeded on low elastic modulus (0.3 kPa) viscoelastic model ECMs showed an increase in cell migration rate and displayed larger cell area, as opposed to those seeded on higher elastic modulus viscoelastic model ECMs.^30^ This contradicts observations on purely elastic model PAH ECMs, where higher elastic modulus model ECMs lead to a higher cell migration rate and a larger projected cell area compared to their more compliant elastic counterparts. Mechanobiology experiments have also investigated fibroblasts. Fibroblasts (MF3) seeded on substrates of similar elasticity, but different loss modulus, displayed significant differences in YAP activation and subsequent proliferation.^40^ YAP nuclear translocation was higher on elastic substrates in comparison to viscoelastic substrates, and was accompanied by higher migration speeds on viscoelastic as opposed to elastic substrates.^41^

Taken together, the examples above clearly illustrate that cells respond differently to elastic and viscoelastic model ECMs, including PAH-based substrates. However, most studies in the literature use PAH substrates with relatively low elastic moduli.^21,30,31,37^ Here, we present a protocol for fabricating model ECMs based on PAH, targeting a wider range of viscoelastic substrates with a fixed storage modulus and tunable loss modulus. Our reported library of tunable viscoelastic model ECMs consists of substrates with storage moduli *G*□ of 3 kPa (*E* ≃ 8 kPa, referred to as soft), 8 kPa (*E* ≃ 25 kPa, referred to as intermediate), and 12 kPa (*E* ≃ 32 kPa, referred to as stiff). For the viscoelastic substrates, the loss moduli *G*□□ were tuned to values of ≃ 300 Pa, 500 Pa, and 700 Pa, for soft, intermediate, and stiff substrates, respectively.^39^ Targeted loss moduli values were achieved by embedding 1.8 % of linear polymer polyacrylamide chains into the soft, intermediate, or stiff elastic PAH networks, adapting previously published protocols.^21,32^ PAHs surfaces were functionalized with collagen type-I to promote cell adhesion. Human lung carcinoma epithelial cells (A549s) were then used to investigate how properties such as cell migration, cell area, and cell adhesion differed between elastic and viscoelastic model ECMs. Overall, we found that the most significant differences exhibited by A549 cells were between stiff elastic and stiff viscoelastic PAH model ECMs. Specifically, cells migrated ∼30% more slowly on elastic model ECMs than on their viscoelastic counterparts. This correlated with larger focal adhesion areas on elastic than on viscoelastic model ECMs. On the other hand, cells migrated at similar rates on soft elastic and soft viscoelastic model ECMs, but focal adhesion areas were around 63% smaller on soft elastic compared to soft viscoelastic. Overall, our findings elucidate the intricate interplay between elastic and viscoelastic properties in regulating epithelial cell mechanobiology, underscoring the significance of time-dependent matrix mechanics in governing epithelial responses. More specifically, our results reveal that epithelial cells distinguish between elasticity and viscoelasticity independent of the storage modulus.

## 2. Materials and methods

### 2.1 Linear elastic polyacrylamide gel preparation

Elastic polyacrylamide hydrogels (PAH) were fabricated following previously reported protocols.^21,32,38,39,41^ Linearly elastic polyacrylamide hydrogels were created by mixing different concentrations of 40% (v/v) acrylamide (Sigma Aldrich, catalog #1610149), and 2% (v/v) bis-acrylamide (Sigma Aldrich, catalog #1610142), leading to 3 different values of the elastic modulus stiffnesses (stiff, intermediate, and soft; see Table 1). Polymerization was initiated by adding 5 μL of 10% w/v ammonium persulfate (APS) (Invitrogen, ref. HC2005), and 0.5 μL of 0.1% (final concentration) of N,N,N,N-tetramethylethylene (TEMED) (Thermo Scientific, Lot # WJ334964), of indicated amounts.^31,38,42^

**Table 1:** Formulations for elastic and viscoelastic polyacrylamide hydrogels (PAH).

|  | Acrylamide (%) | Bis-acrylamide (%) | Linear Acrylamide (%) | Acrylamide from 40% stock solution (mL) | Bis-acrylamide from 2% stock solution (mL) | Linear Acrylamide (mL) | Water (mL) |
| --- | --- | --- | --- | --- | --- | --- | --- |
| Soft E | 5 | 0.30 | 0 | 0.125 | 0.150 | 0 | 0.725 |
| Soft VE | 5 | 0.30 | 1.8 | 0.125 | 0.150 | 0.450 | 0.275 |
| Intermediate E | 8 | 0.25 | 0 | 0.200 | 0.100 | 0 | 0.700 |
| Intermediate VE | 8 | 0.25 | 1.8 | 0.200 | 0.100 | 0.450 | 0.250 |
| Stiff E | 8 | 0.48 | 0 | 0.200 | 0.240 | 0 | 0.560 |
| Stiff VE | 8 | 0.48 | 1.84 | 0.200 | 0.240 | 0.460 | 0.100 |

### 2.2 Linear acrylamide

Linear acrylamide was fabricated by mixing different amounts of 40% acrylamide, milliQ water, APS, and TEMED as summarized in Table 2. Samples were polymerized overnight at 37°C in the dark.^21,31,32,42^ The shear viscosity of the resulting polymer solution was measured with steady shear experimentson an Anton Paar 302e rheometer using a parallel-plate attachment (PP-25 mm). The average zero-shear viscosity, *η*_0_, measured regularly every week over a 5-week period, was 15420 ± 386 mPas, Figure S1. The consistency of the measured values of *η*_0_ suggests that the linear acrylamide chains remained stable over the five-week period, with no signs of degradation. Based on these observations, linear acrylamide solutions were stored for up to 5 weeks and used to fabricate viscoelastic polyacrylamide hydrogels (PAH), as described below.

**Table 2:** Recipe for Linear Acrylamide.

| 40% Acrylamide (mL) | Water (mL) | TEMED (mL) | APS (mL) | NHS 4% |
| --- | --- | --- | --- | --- |
| 1.25 | 8.72 | 0.005 | 0.024 | 0 |

### 2.3 Fabrication of viscoelastic polyacrylamide gels

Viscoelastic PAHs were fabricated by modifying previously described linearly elastic PAH protocols,^21,31,32,42^ by adding the linear acrylamide solution and adjusting the water content as summarized in Table 1. To remove bubbles and minimize dissolved oxygen content, the solution was degassed for 10 minutes, after which TEMED and APS were added. After polymerization, PAHs were fully immersed in PBS and allowed to swell overnight at 4 °C.

### 2.4 Treatment of glass-bottom dishes

Glass-bottom dishes were used as substrates for fabricating elastic and viscoelastic PAHs. The glass portion of the glass-bottom dishes was cleaned with 0.1 M NaOH and allowed to dry overnight. Clean dishes were then treated with (3-aminopropyl)trimethoxysilane, 97% (APTMS) (ThermoFisher Scientific, A11284) for 6 minutes, washed three times with milliQ water (18.2 MΩ·cm at 25 C). Next, glass-bottom dishes were treated with 200 ⍰L of 0.5% glutaraldehyde in PBS (ThermoFisher Scientific, A17876) for 30 minutes, followed by three washes with milliQ water. Samples were dried before further addition of 30 ⍰L of premixed PAH solution. The droplet was subsequently sandwiched between the activated glass well and a clean coverslip (MercedesScientific, 12 mm round #1, #MER R0012), flipped immediately to ensure a horizontal substrate, and allowed to polymerize for 15 minutes. Finally, gels were fully swollen by adding PBS overnight, resulting in PAHs of ∼150 ⍰m in thickness. This protocol has been described in detail elsewhere.^4,21,31^

### 2.5 Preparation of PAH surfaces for cell attachment

To promote cell attachment to both PAHs types (elastic and viscoelastic), the surfaces were functionalized with an adhesive ligand, collagen type I (Col-I). First, the coverslips used in the previous step to create a uniform surface were carefully removed. The surfaces of both elastic and viscoelastic PAH substrates were first activated with Sulfo-SANPAH (ThermoFisher Scientific, A35395). For that purpose, 1 mg of Sulfo-SANPAH was initially dissolved in 200 ⍰L DMSO, yielding a 10 mM solution. Next, Sulfo-SANPAH was further diluted by adding 50 µL of stock solution to 950 µL of HEPES buffer. The diluted Sulfo-SANPAH solution was added to the PAH sample and exposed to UV light for 15 minutes. Samples were rinsed three times with HEPES buffer. The process was repeated once more. Then, 100 ⍰L of Col-I at 100 ⍰g/mL was added to the Sulfo-SANPAH-treated PAH and incubated overnight at 4 C. To conclude, PAH samples were rinsed with PBS and UV-sterilized for 10 minutes before cells were seeded as described below.

### 2.7 Rheology

Elastic and viscoelastic PAHs were characterized using shear rheology, and both strain and frequency dependence were probed. Measurements were performed using an MCR-302e rheometer (Anton Paar) at 25 °C with a sandblasted parallel-plate geometry (PP-25/S, 25 mm diameter). Pre-mixed polyacrylamide solutions were prepared using the monomer acrylamide and the cross-linker bis-acrylamide, as shown in Table 1. The bottom plate was loaded with 510 μL of pre-mixed solution. Next, the sandblasted top plate was slowly lowered until a 1 mm gap was achieved, ensuring that the droplet contacted the top plate and filled the gap. Water was then added to the Peltier-controlled temperature hood (H-PTD220) to prevent the sample from drying. Gels were left to polymerize for 30 minutes. After polymerization, the gap size was reduced slightly to 0.990 mm, resulting in a 1% compressive strain. The storage (*G*□) and loss (*G*□□) moduli were then systematically measured as a function of angular frequency (*ω*) and as a function of shear strain (*γ*). For the angular frequency sweep tests, the frequency *ω* was varied between 0.1-200 rad/s (or equivalently 0.016-31.8 Hz) at constant shear strain (*γ* = 1%), within the linear regime of previously reported elastic PAH measurements.^39^ For shear strain sweep experiments, the shear *γ* was varied between 0.1-50% at constant angular frequency (*ω* = 6.28 rad/s, or 1 Hz). All reported data are presented as the mean ± standard error of the mean calculated from 3 independent tests, each with a freshly prepared and loaded sample.

### 2.9 Cell culture

Adenocarcinoma human alveolar basal epithelial cells (A549) were purchased from Berkeley Biosciences Divisional Services, UC Berkeley. Cells were cultured in growth media composed of Dulbecco’s Modified Eagle Medium-high glucose (DMEM 1X) (ThermoFisher Scientific, 2906246) supplemented with 10% v/v fetal bovine serum (FBS) (Corning, 35-015-CV) in a 100 mm petri dish. No antibiotics were used, as these have been shown to induce metabolic changes.^43^

### 2.10 Single-cell migration time-lapse studies

A549s between passages 2-20 were seeded at 1000 cells/cm^2^ onto sterilized PAH elastic or viscoelastic samples and cultured for 12 hours before imaging, as described below, was initiated. Images were taken every 15 minutes for a total period of 24 hours. We used an inverted epifluorescence microscope (Olympus IX 83 P2ZF) equipped with autofocus and a sterile environmental chamber with temperature (37 °C), humidity, and CO_2_ (5%) control. An Olympus LWD UPLAN FLUOR 20X PH air objective with a numerical aperture (NA) of 0.45 and a working distance of 2.10 mm was used to acquire images in *.tiff format. The static images obtained were then stacked to generate time-lapse movies using ImageJ (Fiji). As soon as the imaging period ended, samples were immediately fixed as described below. Fixed samples were further used for immunostaining and immunofluorescence imaging to quantify focal adhesion sizes.

To analyze the time-lapse movies and to characterize cell migration quantitatively, we curated the data as follows. Cell trajectories that fulfilled the following criteria were used in the evaluation of time dependent displacements: (1) only imaged cells that migrated at least a distance typical of its diameter (∼30 - 40 µm) within the first hour were considered; (2) cell trajectories of cells that contacted other cells were not taken into account; (3) cells that exited the field of view during the imaging process (24 hours) were not considered, and (4) cells that underwent division within the 24-hour period were not considered.

### 2.11 Immunofluorescence imaging

Cells were fixed immediately after the 24-hour time-lapse imaging process was completed. To fix the cells, 200 ⍰L of 4% paraformaldehyde (PFA) (Spectrum, P1010) in PBS was applied for 10 minutes at room temperature (25 °C), followed by three thorough washes with PBS. Fixed cells were then permeabilized with 200 ⍰L of 0.1% Triton X-100 (Sigma, X100) in PBS for 15 minutes and incubated in blocking buffer (2% bovine serum albumin [BSA; Fisher Scientific, BP671-10] in PBS). 4’,6-diamidino-2-phenylindole, dihydrochloride (DAPI) (ThermoFisher Scientific, 62247), Alexa Fluor 488 phalloidin (ThermoFisher Scientific, A12379), and Alexa Fluor 647 anti-paxillin (Santa Cruz Biotechnology, sc-365379) staining solutions were prepared following the manufacturer’s instructions. First, cells were immersed in DAPI solution for 10 minutes, followed by a thorough PBS wash. Then, the cells were immersed in 488 phalloidin for 30 minutes, followed by a thorough wash in PBS. The final staining consisted of immersing cells in 647 anti-paxillin and left overnight at 4°C, followed by a thorough PBS wash. Samples were dried, Prolong Live Antifade Reagent (ThermoFisher Scientific, P36975) was added, and capped with a coverslip to increase fluorescence signal stability. Immunofluorescence images were taken on an LSM 880 upright confocal microscope using a Plan-Apochromat 63x/1.4 Oil DIC M27 objective, resulting in a pixel-to-micron ratio of 0.132. All images were taken at the basal cellular plane and were analyzed with ImageJ FIJI software^45^ following previously reported protocols.^46^

### 2.12 Quantification of focal adhesion areas

Focal adhesion areas were quantified from cell images using ImageJ FIJI.^46^ All images were first converted to 8-bit images. Background subtraction was performed by setting the rolling ball radius parameter to 50 pixels and selecting the sliding paraboloid function of FIJI. Next, the contrast-limited adaptive histogram equalization (CLAHE) function with a block size of 19 was applied to the adjusted images; we set the histogram to 256 and used a maximum slope of 6. The image processing protocol ended with the application of the exponential operator to further minimize background noise effects and adjust brightness. Each image was then manually cropped to exclude everything except the focal adhesions of the cell being analyzed. To quantify focal adhesion areas, the particle plug-in was used, setting a particle size threshold ranging from 0.10 ⍰m^2^ to 15 ⍰m^2^ and a circularity between 0.00 and 0.99.

### 2.13 Statistical Analysis

All statistical analyses were performed using IBM SPSS Statistics and GraphPad Prism. A Student’s t-test was conducted to determine whether differences in elastic and viscoelastic biophysical parameters among the soft, intermediate, and stiff means, with standard errors of the means (SEMs), were statistically significant. (NS = non-significant, * *p* < 0.05, and *** *p* < 0.001).

### 2.14 Cell tracer tool

Areas of migratory cells were also quantified using Marker Tracker. Details of the in-house software are available at https://github.com/MECHANO3B-I-OLOGY/Marker-Tracker. A custom GUI-based application enabled manual delineation of cell boundaries. Users traced contours frame by frame using a brush interface, with optional preprocessing to enhance contrast. From each mask, the software computed the area, perimeter, and centroid using standard contour-based image analysis. Results were exported as structured *.csv files, with optional conversion to physical units based on user-defined pixel scaling. Visual overlays and mask stacks (TIFF format) supported trace validation and reproducibility.

### 2.15 Cell migration quantification via Marker Tracker

Cell migration data analysis was performed using in-house Python-based software: Marker Tracker XYZ (https://github.com/MECHANO3B-I-OLOGY). The Marker Tracker tool recorded the *x-* and *y*-coordinates of selected objects over time. Cells were tracked using an adjustable bounding box (bbox) to ensure the tracker captured the appropriate region of interest (ROI) around the cell. Marker Tracker used the Channel and Spatial Reliability Tracker (CSRT) Python package, which is optimized for tracking deformable objects, such as cells. The centroid coordinates, (*<u>c</u>*_x_, *c*_y_), in the imaging *(xy*) plane for each cell (with identified area) at each point in time were computed using image moments as follows:

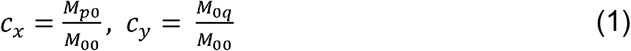

where *M*_*00*_ represents the zero-order moment:

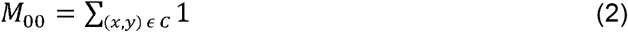

and spatial moments *M*_pq_ are defined by:

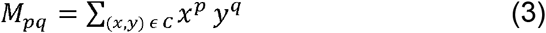

In equations (2) and (3), *C* denotes the contour of the cell. Variables *x, y* within the summation in equation (3) are considered only for locations (points) within the cell contour. This protocol enabled accurate spatial tracking of the cell centroid, even as the moving cell changed shape.

Cell speed was then calculated from the estimated centroid positions by calculating cell (centroid) displacements between image frames. The time increments were determined by the acquisition frame rate. The change in cell centroid position between successive frames was then used to calculate the cell speed. Specifically, the instantaneous speed at time *t* was computed using the equation

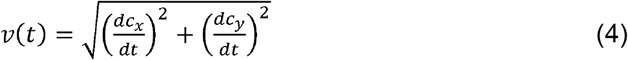

where *c*_x_ and *c*_*y*_ represent the tracked (x,y) coordinates of the cell centroid. The identification of the cell contour *C*, and the cell centroid position (shown in red) is illustrated in Figure S2.

### 2.16 Calculations of the mean square displacement (MSD)

Cells that satisfied the criteria listed above were considered for analysis. For each tracked cell, the location of the cell-centroid in the imaging plane (defined as the x-y plane) was calculated from the images. The mean square displacement (MSD) was then obtained by using the time-sequence of centroid coordinates^44^:

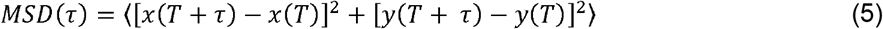

where *x* and *y* represent the centroid coordinates at experimental time *τ*, and *T* is the initial time. To estimate if calculated cell MSDs followed diffusive, sub- or super-diffusive behavior, we calculated the power-law exponent *α*, by fitting MSD data to the following relationship:

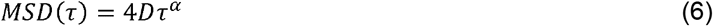

The constant *D* can be identified with the cell diffusion coefficient when *α* = 1. More generally, we treated *D* as a prefactor and used the power-law exponent *α* to quantify the type of migratory behavior. When *α* < 1, cell migration was sub-diffusive; *α* > 1 suggested super-diffusive migration, while *α* = 1 was indicative of Brownian-like diffusion.^44^ The mean square displacement was calculated for each cell based on the trajectory of its centroid. Averages were calculated by combining results from an ensemble of cell trajectories.

## 3. Results

### 3.1 Rheology revealed elastic and viscoelastic tunability of polyacrylamide-based model ECMs

We conducted shear rheology measurements to characterize the shear modulus *G*′ and loss modulus *G*′′ of the linear elastic and viscoelastic model ECMs fabricated. This characterization was done as a function of angular frequency, as well as a function of shear strain to get a complete characterization of the linear viscoelastic properties of the ECM models.

Viscoelastic model ECMs were fabricated by adding linear polyacrylamide chains to the elastic networks, as described in the Materials and Methods section. Figure 1a shows a schematic of the PAH network for elastic and viscoelastic ECM models and illustrates how the addition of linear polyacrylamide chains enhances dissipative effects, thereby allowing tunability of the loss modulus. Following the rheological characterization of the model ECM’s, we evaluated the low-strain and low-frequency values of the storage (*G*′) and loss (*G*′′) moduli.

**Figure 1.**
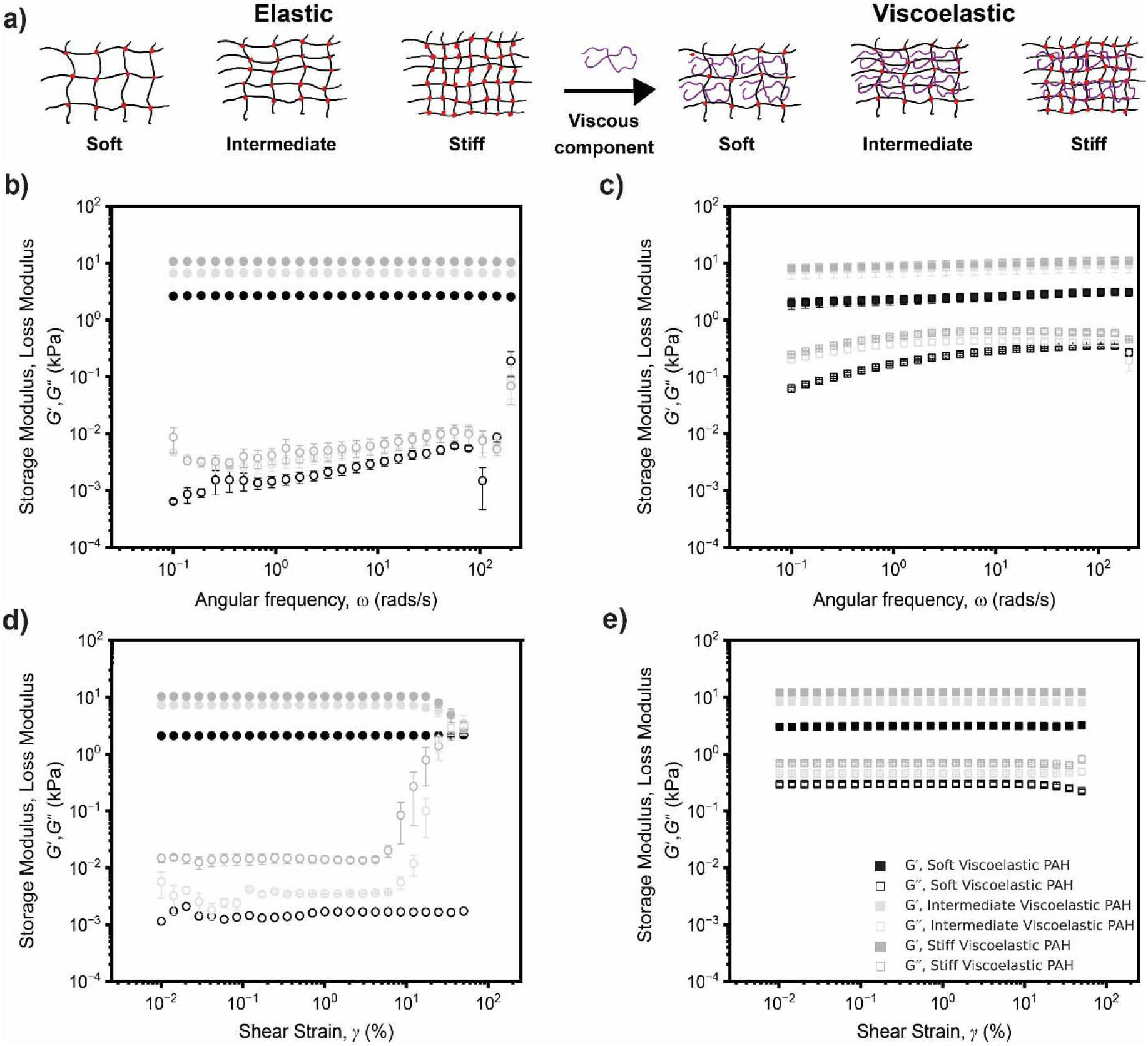
(*a*) Schematic illustrating elastic and viscoelastic PAH model ECM networks. Viscoelastic model ECMs were fabricated by adding linear polyacrylamide chains to the elastic network. (b) Rheological properties of the model ECMs as a function of angular frequency at a shear strain of 1%. We show the storage modulus (*G*□, filled symbols) and loss modulus (*G*□□, open symbols) of model (c) elastic and (d) viscoelastic ECMs, respectively. Rheological properties as a function of shear strain at constant angular frequency of 6.28 rad/s (or 1 Hz). We show the storage modulus (*G*□, filled symbols) and loss modulus (*G*□□, open symbols) for model (e) elastic and (f) viscoelastic ECMs, respectively. Three independent and freshly prepared samples reported. Symbols denote the mean; vertical bars indicate the standard error of the mean of 3 independent measurements for each test.

The storage modulus is controlled by the elasticity of the cross-linked network. In our case, the storage modulus curves attained a non-zero, constant strain-dependent plateau at low frequencies, as is expected from the elasticity of the permanently networked structure. The resting elastic modulus was calculated as the zero-strain limit of this plateau regime. For ideal networks, the loss modulus is expected to vanish under static conditions since there is no viscous dissipation in the absence of flow. Therefore, strictly speaking, at zero frequency, ideal cross-linked viscoelastic substrates respond as purely elastic materials with *G*′′ → 0. Typically, hydrogels may exhibit viscous losses at very low frequencies (with *G*′′ > 0) due to the effects of dangling chains, chain friction, and dissipative effects as water flows through the background network. Here, to enable consistent quantification of linear viscoelastic values, we estimated the zero-strain limit of the storage modulus via extrapolation. These were then compared to the value of the storage modulus obtained from frequency-sweep measurements extrapolated to zero frequency. For the loss modulus, the zero-strain estimate was compared with the low angular frequency (∼0.1 rad/s) value extrapolated from the frequency-sweep measurements.

Figure 1b shows the log-log plots of *G*′ and *G*′′ of elastic soft, intermediate, and stiff model ECMs as a function of angular frequency *ω*. The zero-angular frequency *G*_*0*_□ values were estimated to be *G*_*0*_□= 2.67 ± 0.10 kPa, *G*_*0*_□= 6.59 ± 0.61, and *G*_*0*_□= 10.65 ± 1.00 kPa for soft, intermediate, and stiff elastic model ECMs, respectively. The low-angular frequency *G*_*0*_□□ values were *G*_*0*_□□ = 0.00 ± 0.00 kPa, *G*_*0*_□□ = 0.00 ± 0.00 kPa, and *G*_*0*_□□ = 0.01 ± 0.01 kPa for soft, intermediate, and stiff elastic model ECMs, respectively. Results revealed that within the investigated angular frequency range of 0.1-200 rad/s (or 0.159-31.8 Hz), the elastic model ECMs responded linearly. This is consistent with previous reports in which a combination of tensile testing and shear rheology demonstrated linear-elastic responses in similar model ECMs.^21,32,38,39,41^ Therefore, we concluded that the low *G*_*0*_□□ values measured do not contribute significantly to the mechanical response, and *G*_*0*_□□ will be considered negligible in this study.

Figure 1c shows the log-log plots of *G*′ and *G*′′ of viscoelastic soft, intermediate, and stiff model ECMs as a function of angular frequency *ω*. The zero-angular frequency *G*_*0*_□ values were estimated to be *G*_*0*_□= 1.96 ± 0.77 kPa, *G*_*0*_□= 7.46 ± 3.84 kPa, and *G*_*0*_□= 8.23 ± 0.36 kPa for soft, intermediate, and stiff viscoelastic model ECMs, respectively. The low angular frequency *G*_*0*_□□ values were meanwhile estimated to be *G*_*0*_□□= 0.06 ± 0.006 kPa, *G*_*0*_□□= 0.20 ± 0.040 kPa, and *G*_*0*_□□= 0.246 ± 0.011 kPa for soft, intermediate, and stiff viscoelastic model ECMs, respectively. Results revealed that within the investigated angular frequency range of 0.1-200 rad/s (or 0.159 - 31.8 Hz), the viscoelastic model ECMs *G*′ response was linear; *G*′′ values gradually increased with increasing angular frequency.

Figure 1d shows the log-log plots of *G*′ and *G*′′ for the elastic soft, intermediate, and stiff model ECMs as a function of shear strain *γ*. The estimated values of *G*_*0*_□ at zero-shear strain were *G*_*0*_□= 2.09 ± 0.18 kPa, *G*_*0*_□= 7.2 ± 0.78 kPa, and *G*_*0*_□= 10.24 ± 1.82 kPa for soft, intermediate, and stiff elastic model ECMs, respectively. The zero-shear strain *G*_*0*_□□ values (examined at a frequency of *ω* = 6.28 rad/s, or 1 Hz) were estimated as *G*_*0*_□□ = 0.001 ± 0.001 kPa, *G*_*0*_□□ = 0.01 ± 0.004, and *G*_*0*_□□ = 0.005 ± 0.005 kPa for soft, intermediate, and stiff elastic model ECMs, respectively. These results revealed that within the investigated shear strain range of 0.01-100%, soft elastic model ECMs responded linearly. Similarly, we found that between 0.01 to 10%, intermediate and stiff ECM exhibited a linear response and began to soften above ∼10% shear strain. The sudden decrease in storage modulus *G*’ could indicate network damage induced by the larger shear strains. As in measurements of elastic PAH frequency sweeps, the *G*_*0*_□□ values measured were finite but low.

Figure 1e shows the log-log plots of *G*′ and *G*′′ of viscoelastic soft, intermediate, and stiff model ECMs as a function of shear strain *γ*. We estimated the zero-shear strain *G*_0_□ values to be *G*_0_□= 3.03 ± 0.67 kPa, *G*_0_□= 8.4 ± 0.9 kPa, *G*_0_□= 12.1 ± 1.9 kPa for soft, intermediate, and stiff viscoelastic model ECMs, respectively. We also estimated zero-shear strain *G*_0_□□ values to be *G*_0_□□= 0.292 ± 0.04 kPa, *G*_0_□□= 0.458 ± 0.03 kPa, and *G*_0_□□= 0.690 ± 0.05 kPa for soft, intermediate, and stiff viscoelastic model ECMs, respectively. Results revealed that within the investigated shear strain range of 0.01-100%, the viscoelastic model ECMs’ *G*′ responded linearly. The *G*_0_□□ values measured were significantly higher than those of the linear-elastic counterparts. Additionally, the *G*□□ remained relatively constant over the shear strain range investigated.

Figures 2a and 2b show the average zero-shear strain storage modulus *G*_0_′ and zero-frequency storage modulus values of elastic and viscoelastic model ECMs, where the exact values have been described previously in this section. Likewise, figures 2c and 2d show the average loss modulus *G*_0_□□ of elastic and viscoelastic model ECMs. We found that the zero-shear loss moduli values (evaluated at angular frequency of *ω* = 6.28 rad/s, or 1 Hz) of elastic model ECMs were within 2% or less from values estimated for the corresponding viscoelastic model ECMs. We propose that the loss moduli of the elastic model ECMs did not significantly contribute to the mechanical responses and were therefore treated as ideal elastic. Taken together, our results suggest that the differences in the zero-shear strain and zero-frequency storage moduli between the elastic and viscoelastic model ECMs are not statistically significant, whereas differences in the loss moduli are statistically significant. Values are also summarized in Table 3 and Table 4.

**Table 3.** Limiting values of the shear modulus and loss modulus, *G*□ and *G*□□ obtained from shear-sweep and frequency-sweep tests on elastic PAH model ECMs.

| **Elastic** | **$G_0'$, shear strain (kPa)** | **$G_0''$, shear strain (kPa)** | **$G_0'$, Frequency sweep (kPa)** | **$G_0''$, Frequency sweep (kPa)** |
| --- | --- | --- | --- | --- |
| Soft E | $2.09 \pm 0.18$ | $0.001 \pm 0.001$ | $2.67 \pm 0.05$ | $0.01 \pm 0.00$ |
| Intermediate E | $7.2 \pm 0.450$ | $0.01 \pm 0.002$ | $6.59 \pm 0.357$ | $0.00 \pm 0.00$ |
| Stiff E | $10.24 \pm 1.05$ | $0.005 \pm 0.003$ | $10.65 \pm 0.578$ | $0.00 \pm 0.00$ |

**Table 4.** Limiting values of shear modulus and loss modulus, *G*□ and *G*□□, obtained from shear-sweep and frequency-sweep tests on viscoelastic PAH model ECMs.

| Viscoelastic | $G_0$, shear strain (kPa) | $G''_0$, shear strain (kPa) | $G_0$, Frequency sweep (kPa) | $G''_0$, Frequency sweep (kPa) |
| --- | --- | --- | --- | --- |
| Soft VE | $3.03 \pm 0.39$ | $0.293 \pm 0.023$ | $1.96 \pm 0.44$ | $0.06 \pm 0.004$ |
| Intermediate VE | $8.44 \pm 0.546$ | $0.458 \pm 0.0184$ | $7.46 \pm 2.22$ | $0.20 \pm 0.023$ |
| Stiff VE | $12.12 \pm 1.13$ | $0.689 \pm 0.027$ | $8.23 \pm 0.212$ | $0.246 \pm 0.006$ |

**Figure 2.**
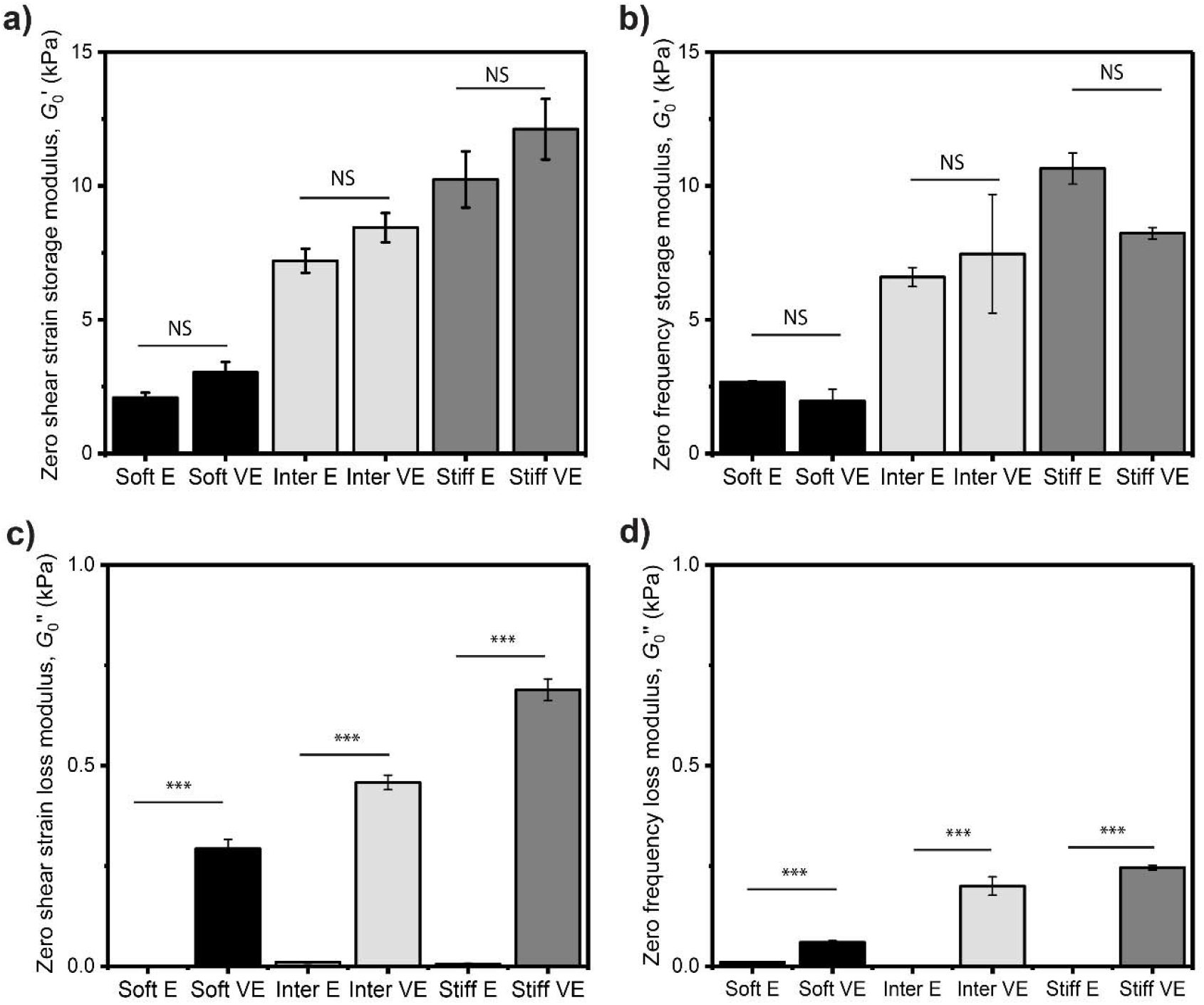
(a) Average zero-strain storage modulus, G_0_′, from shear strain sweep test for soft, intermediate, and stiff elastic and viscoelastic model ECMs, respectively. Experiments were conducted at an angular frequency of 6.28 rad/s (or 1 Hz) (b) Average zero-frequency storage modulus, G_0_′, from angular frequency sweep tests for soft, intermediate, and stiff elastic and viscoelastic model ECMs, respectively. Experiments were conducted at a low strain of 1%. (c) Average estimated zero-strain values of the loss modulus, G_0_′′, from shear strain sweep tests for soft, intermediate, and stiff elastic and viscoelastic model ECMs, respectively. (d) Average low-frequency loss modulus values (for 0.1 rad/s) estimated from angular frequency sweep tests, G_0_′′, for soft, intermediate, and stiff elastic and viscoelastic model ECMs, respectively. An independent t-test was used to assess whether differences between elastic and viscoelastic model ECMs were statistically significant. NS - not significant, * p < 0.05, ** p < 0.01, *** p < 0.001. N = 3 gels per condition.

In addition to frequency- and strain-sweep experiments to estimate the loss and storage moduli from the shear response, we obtained a complementary metric of the time-dependent mechanical response of the ECMs. Specifically, we characterized the relaxation behavior of soft, intermediate, and stiff elastic and viscoelastic model ECMs, as shown in Figure S3 and summarized in Tables S1 and S2. Relaxation data were fitted to expressions with a single relaxation timescale to approximate the dominant dynamical response.

### 3.2 Elasticity and viscoelasticity of PAH model ECMs of similar rigidity alter epithelial migratory behavior

As described in the methods section, we conducted 24-hour time-lapse microscopy studies to monitor and quantify the migratory behavior of A549 cells on elastic and viscoelastic model ECMs. We measured the mean square displacement (*MSD*), the cell motility (migration) exponent (*α*), and the instantaneous cell speed (*V*). For clarity, Figures 3a and 3b show representative cell MSD curves for one A549 cell migrating on collagen type-I-coated soft, intermediate, and stiff elastic and viscoelastic ECMs over a total period of 24 hours. Figure S4 shows *MSD* curves of all cells recorded and for all investigated model ECMs. The slope of the log-log *MSD* versus lag time *τ* was used to extract the cell motility exponent, *α*, which quantifies migration behavior. When *α* = 1, the cell migration mimics pure diffusive behavior. When *α* is < 1, cell migration is sub-diffusive. Finally, when *α* > 1, cell migration behavior is considered super-diffusive; this typically happens due to periods where cells exhibit persistent or directional cell motion. We extracted *α* values by fitting data to Eqn. (6) for specified periods as described next (see SI for details). The MSD data were separated and analyzed into two time periods, 0-10 hours and 10-24 hours, based on an apparent migratory change occurring approximately at 10 hours.

**Figure 3.**
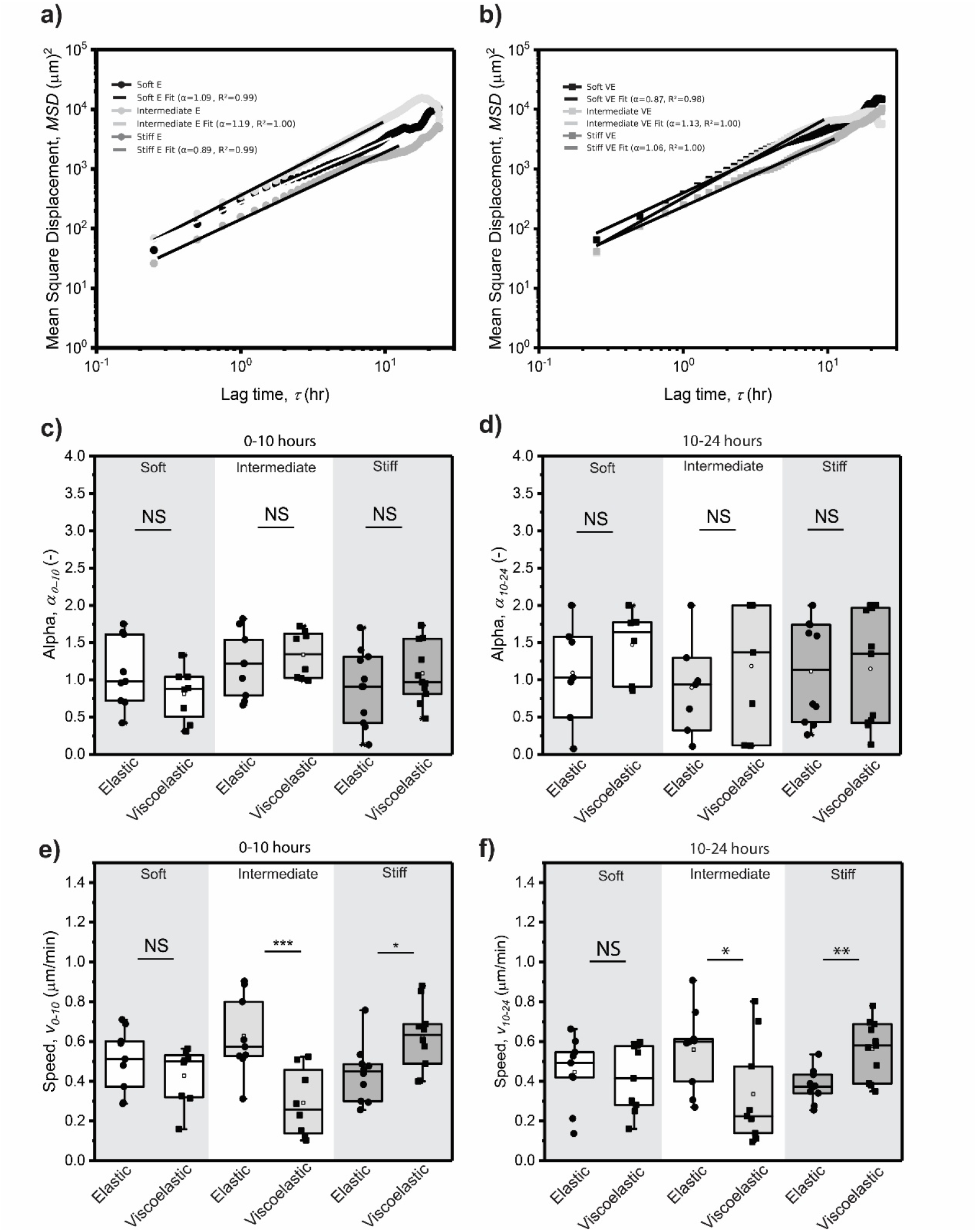
(a) Mean square displacement (MSD) curves versus lag time, *τ*, for A549 cells on soft, intermediate, and stiff elastic model ECMs over 24 hours, respectively. (b) MSD curves versus lag time for A549 cells on soft, intermediate, and stiff viscoelastic model ECMs over 24 hours, respectively. (c) Average A549 cell motility exponent, α, on soft, intermediate, and stiff elastic and viscoelastic model ECMs over 0 to 10 hours. (d) A549 cell migration speed on soft, intermediate, and stiff elastic and viscoelastic model ECMs from 0 to 10 hours, respectively. (e) Average A549 cell motility exponent, α, on soft, intermediate, and stiff elastic and viscoelastic model ECMs over 10 to 24 hours. (f) A549 cell migration speed on soft, intermediate, and stiff elastic and viscoelastic model ECMs from 10 to 24 hours, respectively. NS - not significant, * p < 0.05, ** p < 0.01, *** p < 0.001. N = 10 cells per condition.

Figure 3c summarizes our estimated *α* values for all model elastic and viscoelastic ECMs calculated for cells between 0 and 10 hrs. Averages were computed by taking the average of the single alpha values obtained from each individual cell for each (time) period: 0-10 hours, and 10-24 hours. Specifically, we found that *α* = 1.09 ± 0.16, *α* = 1.19 ± 0.15, and *α* = 0.89 ± 0.16 for elastic soft, intermediate, and stiff model ECMs, respectively. Correspondingly, we found that *α* = 0.87 ± 0.12, *α* = 1.33 ± 0.15, and α = 1.08 ± 0.18 for viscoelastic soft, intermediate, and stiff model ECMs, respectively. Within each substrate type (within each stiffness), we found that the differences in *α* values between elastic and viscoelastic ECMs were not statistically significant using Student’s *t*-tests.

Data for the 10-24 hour period complements the 0-10 hour period data. Figure 3d summarizes our estimates of the *α* values for all model elastic and viscoelastic ECMs calculated for cells between 10 and 24 hrs. We found *α* = 1.09 ± 0.25, *α* = 0.89 ± 0.24, and *α* = 1.11 ± 0.21 for elastic soft, intermediate, and stiff model ECMs, respectively. The *α* values were *α* = 1.26 ± 0.26, *α* = 1.18 ± 0.33, and α = 1.14 ± 0.23 for viscoelastic soft, intermediate, and stiff model ECMs, respectively. Within each substrate type (stiffness), we found that the differences in *α* values between elastic and viscoelastic ECMs were not statistically significant using Student’s *t*-tests.

Figure 3e shows average speeds, *v*, of A549 cells migrating on collagen type-I coated elastic and viscoelastic model ECMs over the 10-hour period. Overall, cells on elastic model ECMs exhibited different migration patterns than those on viscoelastic counterparts. Cells on soft, elastic ECMs migrated faster, with higher instantaneous speeds, than those on stiff, elastic ECMs. Interestingly, cells on intermediate elastic ECMs migrated faster than cells on both soft and stiff ECMs. The average speeds were *v* = 0.5 ± 0.1 ⍰m/min, *v* = 0.6 ± 0.1 ⍰m/min, and *v* = 0.4 ± 0.1 ⍰m/min for cells migrating on soft, intermediate, and stiff elastic model ECMs, respectively. Cells on soft viscoelastic model ECMs migrated at similar speeds as cells on intermediate viscoelastic model ECMs. However, cells on stiff viscoelastic model ECMs migrated faster than on both soft and intermediate model ECMs. We found *v* = 0.4 ± 0.1 ⍰m/min, *v* = 0.3 ± 0.1 ⍰m/min, and *v* = 0.6 ± 0.1 ⍰m/min for soft, intermediate, and stiff viscoelastic, respectively. In summary, cells on intermediate viscoelastic model ECMs migrated 54% slower than on their elastic counterparts, and cells on stiff elastic model ECMs migrated 29% slower than in their viscoelastic counterparts.

Figure 3f shows the average speed of A549 cells migrating on collagen type-I coated elastic and viscoelastic model ECMs for the period from 10-24 hours. Overall, cells on elastic model ECMs exhibited different migration patterns than those on their viscoelastic counterparts. In particular, cells on soft elastic ECMs migrated faster compared to those on stiff elastic ECMs. Interestingly, cells on intermediate elastic ECMs migrated faster in comparison to both soft and stiff ECMs. We estimated *v* = 0.5 ± 0.1 µm/min, *v* = 0.6 ± 0.1 µm/min, and *v* = 0.4 ± 0.0 µm/min for cells migrating on soft, intermediate, and stiff elastic model ECMs, respectively. Cells on soft viscoelastic model ECMs migrated at similar speeds to cells on intermediate viscoelastic model ECMs. However, cells on stiff viscoelastic model ECMs migrated faster than cells on both soft and intermediate model ECMs. We estimated *v* = 0.4 ± 0.1 µm/min, *v* = 0.3 ± 0.1 µm/min, and *v* = 0.6 ± 0.1 µm/min for cells migrating on soft, intermediate, and stiff viscoelastic, respectively. In summary, our experiments indicate that cells on intermediate viscoelastic model ECMs migrated 41% more slowly than on their elastic counterparts, and cells on stiff elastic model ECMs migrated 32% more slowly than on their viscoelastic counterparts.

To assess whether cell speeds remained similar or changed dramatically over the 24-hour observation period, average cell speeds were calculated for 4 intervals: 0-6 hours, 6-12 hours, 12-18 hours, and 18-24 hours. Figures S5 and S6 show the speeds of cells recorded on all investigated model ECMs. Overall, A549 cell speed on soft, intermediate, and stiff elastic and viscoelastic model ECMs varied with time, as evidenced by differences across the 4 time periods. Cells migrating on elastic and viscoelastic soft model ECMs migrated at similar speeds. Similar behaviors were observed in cells migrating on elastic and viscoelastic stiff model ECMs, that is, migrated at similar speeds. However, this was not the case for cells migrating on elastic and viscoelastic intermediate-model ECMs. Cells on the elastic intermediate model ECMs migrated faster. Overall, these findings indicate that while ECM stiffness generally drives comparable migration speeds on soft and stiff elastic and viscoelastic substrates, intermediate stiffness uniquely reveals a pronounced dependence on ECM mechanics.

### 3.3 Projected cell areas of migratory cells are similar on elastic and viscoelastic PAH model ECMs

To study the impact of the loss modulus on cell motility and cell-substrate mechanics, we investigated time-dependent projected cell areas as cells moved on the model ECMs. For each frame imaged, an instantaneous cell area was determined as described in the methods section. Figures 4a and 4b show the average projected cell area, *A*, for cells on soft, intermediate, and stiff elastic and viscoelastic model ECMs. We observe that cells moving on elastic model ECMs reached homeostatic behavior after ∼10 hours and followed the expected trend of increasing area with increasing stiffness. Cells on viscoelastic model ECMs reached homeostatic behavior faster, after approximately 7 hours.

**Figure 4.**
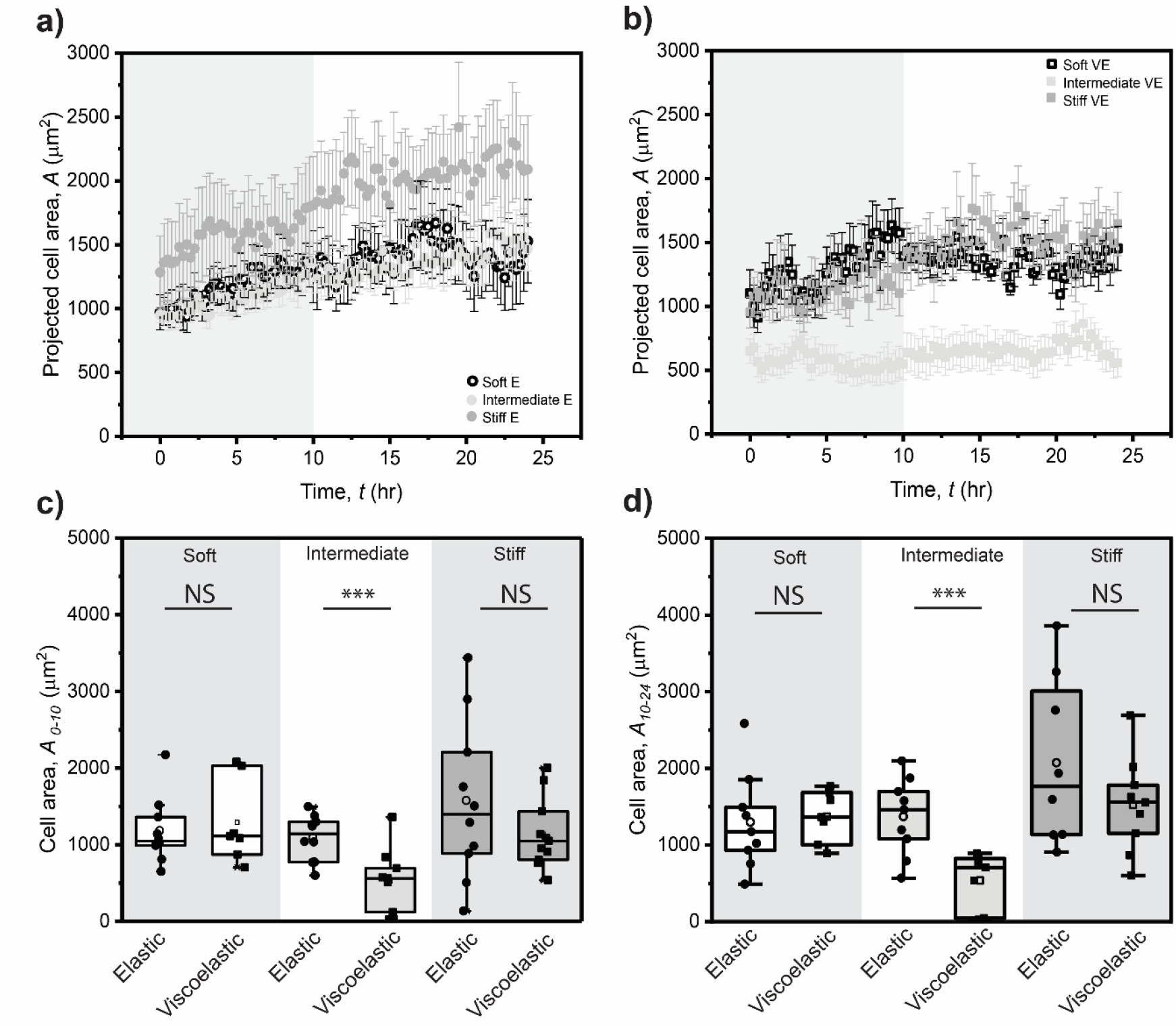
(a) A549 cells projected cell area on soft, intermediate, and stiff elastic model ECMs over 24 hours, respectively. (b) A549 cells projected cell area on soft, intermediate, and stiff viscoelastic model ECMs over 24 hours, respectively. (c) Cell area on soft, intermediate, and stiff elastic and viscoelastic model ECMs from 10 – 24 hours. (d) Cell area on soft, intermediate, and stiff elastic and viscoelastic model ECMs after 24 hours. An independent t-test was used to assess whether differences in cell behavior between elastic and viscoelastic model ECMs were statistically significant. NS - not significant, * p < 0.05, ** p < 0.01, *** p < 0.001. N = 10 cells per condition.

Interestingly, cells on intermediate viscoelastic model ECMs had smaller projected cell areas than on soft and stiff viscoelastic model ECMs, whereas they exhibited similar projected cell areas on soft and stiff viscoelastic ECMs. In contrast, cells exhibited a larger projected cell area on intermediate elastic ECMs than on viscoelastic ECMs. Finally, cells had a larger cell area on stiff elastic ECMs than on stiff viscoelastic ECMs.

Figure 4c and 4d show the average cell area between 10 hours and 24 hours (*i*.*e*., homeostasis/steady state) and the average cell area after 24 hours for soft, intermediate, and stiff elastic and viscoelastic model ECMs. Averages were calculated from 10 cells per condition. Figure 4c shows the average cell area from 10 hours to 24 hours for cells on elastic and viscoelastic model ECMs. The average measured cell areas were 1298.3 ± 210.3 µm^2^, 1371.4 ± 167.0 µm^2^, and 2072.4 ± 387.0 µm^2^ for elastic soft, intermediate, and stiff model ECMs, respectively. Similarly, the average measured cell areas were 1370.0 ± 126.5 µm^2^, 537.9 ± 135.0 µm^2^, 1522.2 ± 208.1 µm^2^ for viscoelastic soft, intermediate, and stiff model ECMs, respectively. Figure 4d shows the final cell area at *t* = 24 hours for cells on elastic and viscoelastic model ECMs. The average measured cell areas were 1527.5 ± 390.1 µm^2^, 1467.4 ± 179.7 µm^2^, and 2089.96 ± 469.5 µm^2^ for elastic soft, intermediate, and stiff model ECMs, respectively. Similarly, the average measured cell areas were 1364.6 ± 192.3 µm^2^, 557.5 ± 154.9 µm^2^, 1524.8 ± 279.7 µm^2^ for viscoelastic soft, intermediate, and stiff model ECMs, respectively. Interestingly, average cell areas at 24 hours on viscoelastic intermediate model ECMs were 62% smaller in comparison to the average for cells on elastic intermediate model ECMs. Lastly, average cell areas on elastic stiff model ECMs were similar to those on viscoelastic stiff model ECMs after 24 hrs.

### 3.4 Focal adhesion size depends on loss modulus for soft and stiff PAH model ECMs, but not for intermediate

To gain insight into the interplay between elasticity and viscoelasticity and focal adhesion complexes, which are involved in the mechanosensory machinery of migratory cells, we quantified paxillin at focal adhesions to estimate focal adhesion sizes in A549 cells on elastic and viscoelastic model ECMs after a 24-hour time-lapse. Figures 5a, 5b, and 5c show representative images of paxillin stained on elastic and viscoelastic soft, intermediate, and stiff model ECMs. Figure 5d summarizes the focal adhesion area, *A*_*FA*_, for soft, intermediate, and stiff elastic and viscoelastic model ECMs. Cells on soft elastic model ECMs exhibited a smaller focal adhesion area compared to cells on intermediate elastic model ECMs. However, cells on stiff elastic model ECMs exhibited larger focal adhesion areas than those on soft and intermediate elastic model ECMs. This trend is expected and has been observed for multiple adherent cell types.^21,30,40^ The average measured focal adhesion areas were 0.3 ± 0.1 µm^2^, 0.6 ± 0.1 µm^2^, 0.8 ± 0.1 µm^2^ for soft, intermediate, and stiff elastic, respectively. Interestingly, cells on stiff viscoelastic ECMs formed smaller focal adhesion areas than on both soft and intermediate viscoelastic ECMs. The average measured focal adhesions were 0.8 ± 0.1 µm^2^, 0.7 ± 0.1 µm^2^, and 0.5 ± 0.1 µm^2^ for soft, intermediate, and stiff viscoelastic, respectively.

**Figure 5.**
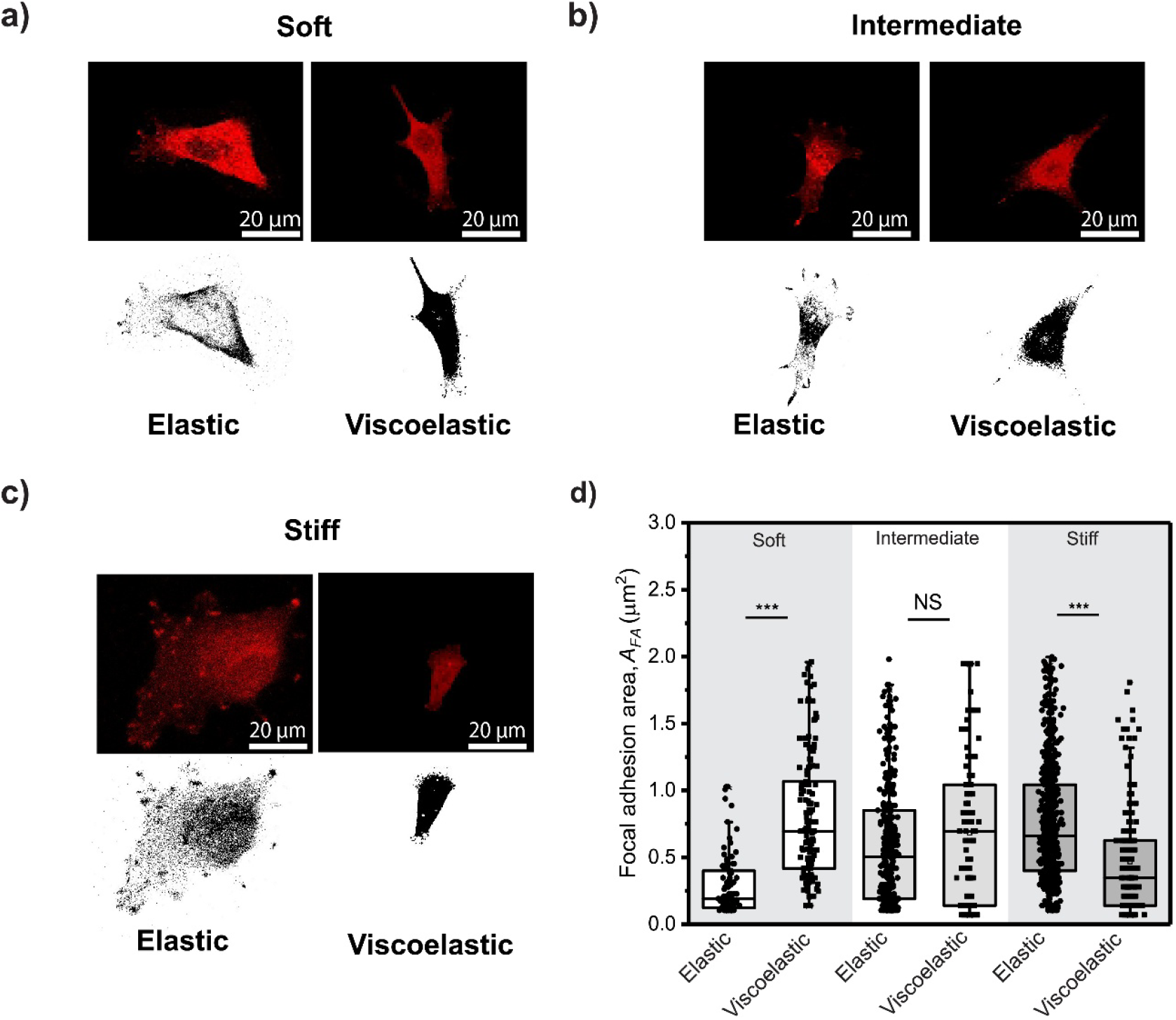
(a) Immunofluorescence image of paxillin in A549 cells on soft elastic and viscoelastic model ECMs. (b) Immunofluorescence image of paxillin in A549 cells on intermediate elastic and viscoelastic model ECMs. (c) Immunofluorescence image of paxillin in A549 cells on stiff elastic and viscoelastic model ECMs. (d) Focal adhesion area of A549 cells on soft, intermediate, and stiff elastic and viscoelastic model ECMs. An independent t-test was used to determine if the differences in cell behavior between elastic and viscoelastic model ECMs were statistically significant. NS - not significant, * p < 0.05, ** p < 0.01, *** p < 0.001.

Finally, we compared the focal adhesion area of cells on elastic model ECMs with those on viscoelastic model ECMs. Cells on soft elastic model ECMs assembled focal adhesion areas that were 65% smaller compared to those on soft viscoelastic model ECMs. However, cells on intermediate elastic model ECMs showed focal adhesion areas comparable to those on intermediate viscoelastic ECMs. Interestingly, cells on stiff elastic model ECMs exhibited a 65% larger focal adhesion area than those on stiff viscoelastic model ECMs.

## 4. Discussion

While PAH based viscoelastic model ECMs have been reported in the literature, currently used substrates span a limited range of stiffness values. Furthermore, the effects of ECM viscoelasticity on migratory cells remain incompletely understood. Therefore, our first objective was to expand the range of mechanical properties (that is increase attainable values in the stiffness-viscoelastic phase space) and thereby increase the library of tunable PAH for mechanobiology studies. The motivation for focusing on PAH hydrogels is, in part, due to the relative ease in tuning PAH’s mechanical and chemical properties,^47^ and the mechanobiology field’s familiarity with PAHs.^48–54^ The protocols we describe and use enable independent control of the storage and loss moduli over a wider range of stiffness than previously reported.^21,25,31,32,37^ Specifically, consistent with previous studies, we tuned the loss modulus by incorporating linear acrylamide polymer chains into elastic PAH networks crosslinked with acrylamide and bisacrylamide. Our fabrication protocols enabled us to cast viscoelastic PAHs with shear moduli up to 12 kPa (equivalent to Young’s modulus of ∼32 kPa), exceeding values typically reported in the literature. While we did not explore higher values of stiffness, the strategies described here can be expanded easily to fabricate very stiff viscoelastic substrates. For the library of PAHs we study, the loss modulus is within ∼10% of the storage modulus, similar to ratios reported for stromal or connective tissues, ^55–58^ making these substrates biomimetically relevant.

We performed several rheological tests to validate the independent tunability of the storage and loss moduli of our model PAH ECMs: shear-strain sweeps, angular-frequency sweeps, and relaxation modulus measurements. Frequency sweep and strain sweep data were used to estimate the loss and storage modulus in the linear viscoelastic limit, valid for low strain and at low frequencies (∼ 0.1-1 rad/s relevant to mechanobiology studies). We observed similar storage moduli between elastic and viscoelastic model ECMs in both shear tests, indicating that differences were not statistically significant. However, the differences in the loss modulus between the elastic and viscoelastic ECMs were statistically significant. The responses were valid for small strains and low frequencies, relevant to timescales of relevant cellular processes, such as focal adhesion turnover,^59–61^ conformational changes of mechanotransducers,^62–64^ or lamellipodial and filopodia formation.^65,66^ The elastic and viscoelastic responses were further confirmed by compression-relaxation tests; as expected, elastic model ECMs exhibited an instantaneous response, whereas viscoelastic model ECMs exhibited a time-dependent response. Notably, the relaxation time for the intermediate viscoelastic model ECMs was significantly longer.

One of the reported advantages of PAH ECMs is their intrinsic optical transparency, which facilitates imaging of cellular processes using inverted optical and epifluorescence microscopes. All our elastic PAH model ECMs have retained this property. This suggests that the linear polyacrylamide chains were evenly dispersed in the elastic network. However, among the viscoelastic PAH model ECMs, only the intermediate-rigidity ECM retained optical transparency. For the soft and stiff viscoelastic PAH model ECMs, nevertheless, the resulting substrates were translucent. UV-Vis absorbance measurements validated this observation, as shown in Figure S7. This is most likely due to a combination of immiscibility at the working concentrations, and microphase separation of the linear polyacrylamide chains during the curing process of the elastic network. Imaging cells through these ∼150 µm-thick translucent substrates (as estimated from Z-stacks) posed challenges for accurately tracing cell boundaries with our computational tools (the Marker tracker tool described in Materials and Methods) and thus required manual tracing. To overcome this optical limitation, upright microscopy can be used, as was the case to image focal adhesions. In summary, these PAH-based model ECMs expand the tunability of the mechanical niche microenvironment for mechanobiology studies by combining complementary imaging modalities.

Our next objective was to evaluate how the loss modulus affected epithelial cell mechanobiology, specifically focusing on cell mean-squared displacement, migration, cell area, and focal adhesion area. Previous studies have shown that cells can differentiate between elastic and viscoelastic model ECMs, and their responses vary depending on the specific cell type.^30,37^ In these studies, cellular responses varied depending on the ECM ligands presented to cells (*e*.*g*., collagen, fibronectin, or laminin), the model ECM crosslinking parameters, and the location of immobilized ECM ligands, either within the elastic network, within embedded linear polymer chains, or both.^21^ A previous study showed that when only linear polymer chains were functionalized with collagen, cells did not adhere; however, when fibronectin was used, cells adhered.^33^ Here, we functionalized the entire surface using the UV-activated crosslinker Sulfo-SANPAH and collagen type I at 100 µg/mL, functionalizing both model ECM components, the elastic network and surface-exposed linear polyacrylamide chains.

It is well known that increased substrate stiffness promotes cell migration, area, and proliferation, including A549s. For example, collective cell migration of A549s was higher on polydimethylsiloxane (PDMS) on substrates with a stiffness of 18.3 MPa than on 1.4 or 3.4 MPa PDMS.^67^ Tissues, however, are viscoelastic, rather than purely elastic, as is the case with PDMS and other model mechanobiology substrates, such as PAHs. Understanding how energy dissipation in soft materials, quantified by the loss modulus or relaxation constants of model ECMs, regulates cellular mechanisms has attracted considerable attention in recent years. Our study focused on single-cell migration of A549s on PAH-based model ECMs with tunable loss modulus. The combination of cell-type and model ECMs suggests that our study can be considered as a relevant model system that can be further extended to investigate ECM mechanics in cancer metastasis. For instance, ECM mechanics influences how adenocarcinoma cells that have undergone an epithelial-to-mesenchymal transition (EMT) migrate and become invasive, a process required for metastasis. Our results show that an increase in the loss modulus affected cell velocity on intermediate and stiff substrates, but not on soft substrates. Furthermore, the responses between intermediate and stiff substrates were of opposite trend, suggesting, combined with previous literature, that there is no universal trend and that potentially different mechanosensory signaling pathways were activated. Our stiff viscoelastic findings are analogous to those observed for non-tumorigenic human MCF10A cells seeded on alginate-based model ECMs, in which cell migration increased on viscoelastic substrates compared with elastic counterparts.^68^

Another study using MCF10A cells as well demonstrated that on extremely soft viscoelastic (*E* ∼0.3 kPa) PAH-based model ECMs, cells migrated faster than on their elastic counterpart, while migration speed decreased on stiff viscoelastic compared to stiff elastic.^30^ In our case, cells migrated at higher speeds on stiff viscoelastic than on stiff elastic, while cells migrated at slower speeds on intermediate viscoelastic than on intermediate elastic. Interestingly, on soft PAH model ECMs, speeds were similar on viscoelastic and elastic substrates. Yet the migration or motility exponent, which quantifies the form of the mean square displacement, changed significantly from soft elastic (α = 1.09) to soft viscoelastic (α = 0.87) ECMs. Cells on soft, elastic surfaces displayed hindered migratory behavior rather than purely random (Brownian-like) diffusive motion. Indeed, cell migration was significantly hindered on intermediate viscoelastic model ECMs (*G*⍰ ∼ 8 kPa, with relaxation times of ∼3 seconds), as seen in Figure S3. However, the *MSD* curves remained relatively similar for cells moving on intermediate elastic (*α* = 1.19) and intermediate viscoelastic (*α* = 1.13) model ECMs. In contrast, cells on stiff model ECMs (*G*⍰ ∼ 12 kPa, with shorter relaxation times of approximately 0.5 seconds) promoted cell migration. As expected, *MSD* curves differ qualitatively for cells on stiff elastic (*α* = 0.89) and on stiff viscoelastic (*α* = 1.06) model ECMs. A possible explanation for this, motivated by motor clutch models and theories for cells migrating on viscoelastic model ECMs,^25,37^ is that cells on intermediate viscoelastic model ECMs may undergo motor clutch dynamics following “load and fail” regimes. This may lead to weaker focal adhesion forces (indicating immaturity), increased retrograde flow, reduced spreading, and subsequently slower migration. In contrast, cells on stiff viscoelastic model ECMs may experience “frictional slippage,” resulting in smaller focal adhesion sizes and shorter adhesion lifetimes, whereas larger focal adhesions form on stiff elastic ECMs, as shown in Figure 5d. Previous studies using fibroblasts (HMF3) have shown similar findings: viscoelastic model ECMs with higher storage modulus and enhanced cell migration.^40^

Matrix elasticity significantly influences cell spreading area; however, results in the literature regarding cell area for cells interacting with viscoelastic model ECMs have been mixed. Previous studies have shown that fibroblasts exhibit a larger cell area on elastic model ECMs (2428.93 ± 864.71 μm^2^) compared to cells seeded on viscoelastic ECMs (1296.73 ± 311.62 μm^2^) and cells seeded on glass (1792.61 ± 487.09 μm^2^).^40^ This decrease in cell area may be due to the cells’ inability to form large and stable focal adhesions. However, some studies have reported a higher spreading area for cells on viscoelastic ECMs than on their elastic counterparts.^37^ To further characterize the behavior of migratory cells, we analyzed their instantaneous projected cell area over 24 hours. We then compared the cell area at the beginning and end of the time-lapse study, as defined in Section 2.10. Our findings showed that the initial and final cell areas of migratory cells were not statistically different on soft or stiff viscoelastic vs elastic model ECMs. However, cells on intermediate model ECMs and on viscoelastic ECMs showed decreased cell spreading (projected cell area) in both the first half (parsed from t = 0 to 10 hours) and the second half (parsed from *t* = 10 to 24 hours) of the time-lapse studies. This decrease in cell area may be due to cells’ inability to form strong or mature focal adhesions, which prevented them from spreading or slipping during migration, as observed in the cell migration data. Similar trends in cell area have been reported for human airway smooth muscle (HASM) and human prostate carcinoma epithelial (22Rv1) cells on soft tunable elastic and viscoelastic model ECMs.

Lastly, we analyzed the focal adhesion area in relation to the elastic and viscoelastic model ECMs to better understand cell migration. Focal adhesion size may predict or correlate with cell migration on elastic substrates.^69^ We observed that cells formed larger focal adhesions as substrate stiffness increased; this trend was not seen for cells on viscoelastic substrates. Some have reported no significant variation of epithelial cell focal adhesion area on what others have referred to as soft (∼ 0.3 kPa) and stiff (∼ 3 kPa) elastic and viscoelastic model ECMs.^30^ While fibroblasts have displayed significant differences in stiff (∼14 kPa) model elastic and viscoelastic ECMs.^40^ In this study, we examined the focal adhesion area 24 hours after cell seeding on elastic and viscoelastic model ECMs. Interestingly, we observed a larger focal adhesion area on soft viscoelastic substrates compared to their elastic counterparts; however, cell migration did not show significant differences. While the focal adhesion area on intermediate model ECMs remained similar, their migratory behavior differed. Finally, cells on stiff viscoelastic model ECMs exhibit smaller focal adhesions, while migration increases compared to their elastic counterparts.

Cell signaling via the underlying mechanical substrate has been demonstrated for cells on substrates that are neither too stiff nor too compliant.^9,14,15^ Recent theoretical studies by us and collaborators show that substrate stiffness, cell migration rates, and cell-substrate stresses affect the frequency of contacts between neighboring cells, and the ability of migratory cells to move persistently .^70,71^ Our experimental findings highlighting the complex interplay between substrate (ECM) elastic and viscoelastic properties in regulating epithelial cell responses strongly suggest that the role of substrate viscoelasticity must also be considered to understand cell-cell interactions, and emergent long-ranged behavior such as durotaxis and adurotaxis.

## 5. Conclusion

We created a tunable PAH-based viscoelastic platform with storage moduli comparable to those of their elastic counterpart to investigate the response of cell mechanobiology to loss moduli. Using Adenocarcinoma lung epithelial cells (A549s) as a model cell line, we evaluate mean-squared displacement, cell migration, cell area, and focal adhesions on both elastic and viscoelastic model ECMs. Our analysis shows that A549 cell migration is enhanced or hindered on model ECMs with storage moduli above ∼3 kPa, depending on the substrate relaxation time. We also observed a significant decrease in focal adhesion size on stiff viscoelastic model ECMs, correlating with an increase in cell migration speed. Our results suggest that viscoelasticity influences cell migration above a certain stiffness value, depending on the relaxation time of the substrate. We also observe that there is no true correlation between cell migration and focal adhesion on a viscoelastic substrate, as traditionally observed on elastic substrates with increasing storage modulus.

## 6. Conflicts of interest

No conflicts of interest to declare

## 7. Acknowledgements

A.M.S., A.G., and R.C.A.E. acknowledge funding from the NSF-CREST: Center for Cellular and Biomolecular Machines through the support of the National Science Foundation (NSF) Grant No. NSF-HRD-1547848. A.M.S and R.C.A.E. acknowledge funding from the Tobacco-Related Disease Research Program through the support of the University of California Office of the President Grant No. T31KT1583 awarded to R.C.A.E. A.M.S., and A.G. acknowledge funding from the CAREER NSF Grant No. CBET 2047210 awarded to A.G., A.M.S. acknowledges funding from the UC Merced Graduate Dean’s Dissertation fellowship.

## 8. Supplementary Information

**Figure S1**. Stability of the viscosity of linear polymer chains

**Figure S2**. Cell centroid for speed calculations

**Figure S3**. Relaxation of soft, intermediate, and stiff elastic and viscoelastic model ECMs

**Table S5**. Relaxation values of elastic model ECMs

**Table S6**. Relaxation values of viscoelastic model ECMs

**Figure S4**. Total mean square displacement of soft, intermediate, and stiff elastic and viscoelastic model ECMs over 24 hours

**Figure S5**. A549 cell migration speed on soft, intermediate, and stiff elastic and viscoelastic model ECMs

**Figure S6**. Average cell speed across different time increments on soft, intermediate, and stiff elastic and viscoelastic model ECMs.

**Figure S*7***. UV-Vis spectra comparison of Polyacrylamide soft, intermediate, and stiff viscoelastic model ECMs

## Notes

### Competing Interest Statement

The authors have declared no competing interest.

### Summary of Updates

Added clarifying statements in the results sections. Expanded methodology in the SI.

