## Supplementary material for "Adenocarcinoma cell mechanobiology is altered by the loss modulus of the surrounding extracellular matrix": S

**Figure S1.** Stability of the viscosity of linear polymer chains

**Figure S2.** Cell centroid for speed calculations

**Figure S3.** Relaxation moduli of soft, intermediate, and stiff elastic and viscoelastic model ECMs estimated from rheometry.

**Table S1.** Relaxation values of elastic model ECMs estimated from rheometry.

**Table S2.** Relaxation values of viscoelastic model ECMs estimated from rheometry.

**Figure S4.** Mean square displacement (MSD) versus lag time for  cells moving on soft, intermediate, and stiff elastic and viscoelastic model ECMs evaluated over a period of 24 hours

**Figure S5**. A549 cell migration speed of cells on soft, intermediate, and stiff elastic and viscoelastic model ECMs

**Figure S6.** Average cell speeds evaluated over different time increments on soft, intermediate, and stiff elastic and viscoelastic model ECMs.

**Figure S7**. Comparison of Polyacrylamide soft, intermediate, and stiff viscoelastic model ECMs using UV-Vis spectra.

### 1. Evaluation of shear viscosity to test for material degradation

To determine the stability of linear polymer chains and determine if degradation occurred over time, shear viscosity tests were performed over a 5-week period. The shear viscosity, η, was measured using the rheometer (Anton Paar MCR-302e , with a parallel plate attachment PP-25/S, 25 mm diameter) by applying shear rates from 0.1 $\frac{1}{s}$ to 100 $\frac{1}{s}$.

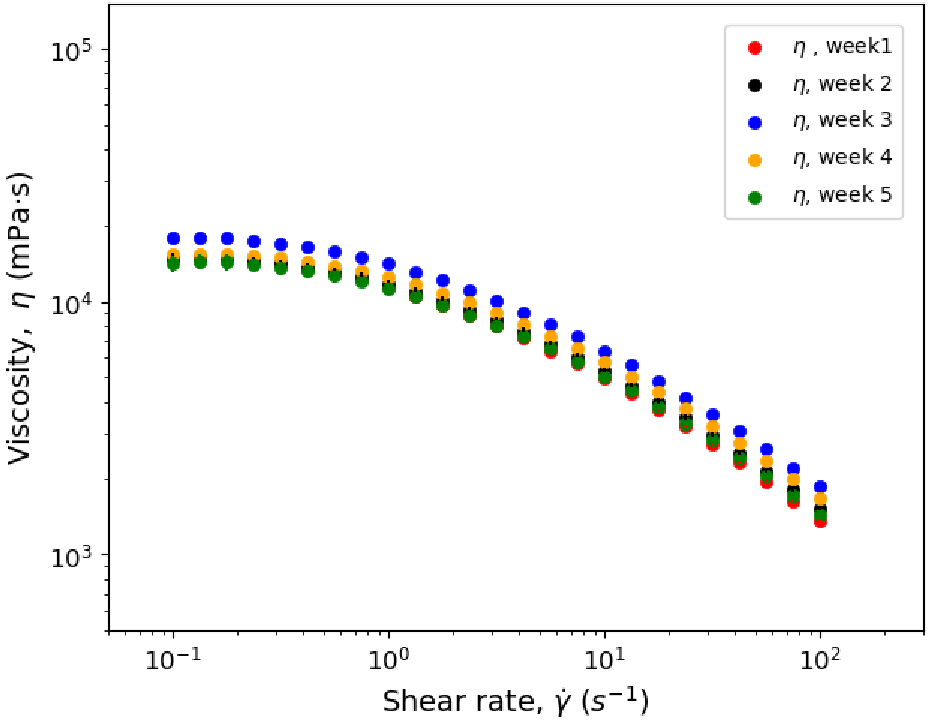


**Figure S1.** Viscosity of linear acrylamide solution as a function of shear rate estimated periodically over a total time of 5 weeks.

### 2. Cell velocity obtained using the cell centroid

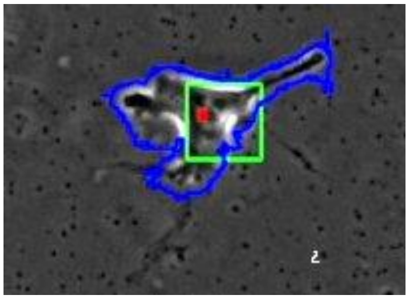

**Figure S2.** A549 cell migration tracked uusing image analysis. The cell centroid (red) and cell boundary (blue) of cell are both obtained from the image.

### 3. Relaxation modulus estimates of elastic and viscoelastic model ECMs

To determine the relaxation behavior of elastic and viscoelastic PAHs, relaxation moduli tests were performed using the rheometer. The relaxation modulus, $G_R(t)$, was measured by applying an instantaneous shear strain, $\gamma$, of 10% (at time $t$ = 0 s) and maintaining it for 1000 s. The characteristic relaxation time, $\tau$, amplitude decay, $A$, and equilibrium shear modulus, $G_\infty$, were extracted using a single exponential decay function:

$$G_R(t) = (G_0 - G_\infty)\, e^{\frac{-t}{\tau}} + G_\infty \qquad \text{(SI 1)}$$

where $t$ denotes time. Fits were performed using Origin Lab version 2017. The form of equation (SI 1) was motivated by simplicity and provides a minimal model with a single dominant relaxation time.

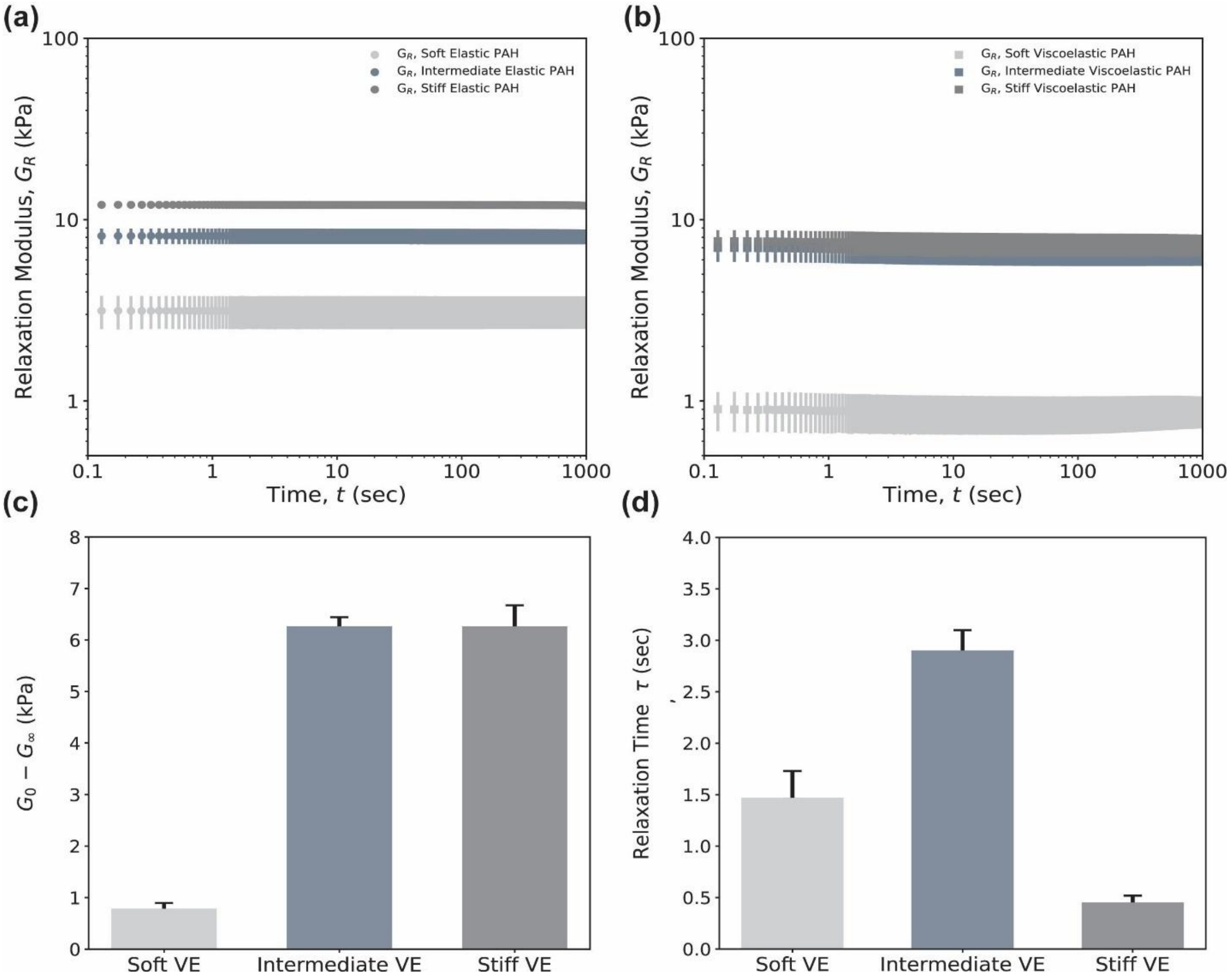


**Figure S3.** (a) and (b) Relaxation modulus after an instantaneous shear strain of 10% as a function of time for model elastic and viscoelastic ECMs, respectively. Three independent and new samples per condition are reported. Shown is the mean and the standard error of the mean of 3 independent measurements for each test. (c) Average shear modulus amplitude difference, $G_0$ - $G_\infty$, for soft, intermediate, and stiff viscoelastic model ECMs, respectively. (d) Average relaxation time, τ, for soft, intermediate, and stiff viscoelastic model ECMs, respectively. Data reported in panels (c) and (d) come from relaxation tests.

As an additional time-dependent mechanical response, we further evaluated the relaxation behavior of soft, intermediate, and stiff elastic (Figure S2a), and viscoelastic (Figure S2b) model

ECMs. Figure SI2a shows the log-log curve of the relaxation modulus, $G_{R,}$ as a function of time. The measured zero relaxation modulus $G_{Ro}$ values at $t_0$ = 1.0 s were $G_{Ro}$ = 3.14 ± 0.65 kPa, $G_{Ro}$ = 8.14 ± 0.81 kPa, and $G_{Ro}$ = 12.1 ± 0.39 kPa for soft, intermediate, and stiff elastic model ECMs, respectively. The measured $G_\infty$ values at $G_\infty$ = 3.14 ± 0.00 kPa, $G_{Ro}$ = 8.07 ± 0.001 kPa, and $G_{Ro}$ = 12.1 ± 0.39 kPa for soft, intermediate, and stiff elastic model ECMs, respectively. Our results reveal that within the investigated time of 1.03-1000 s, elastic model ECMs did not display any relaxation behavior.

Figure S2b shows the relaxation modulus, $G_{R,}$ values as a function of time plotted in log-log form. The measured zero relaxation modulus $G_{Ro}$ values at $t_0$ = 1.0 seconds (when samples begin to relax) were 0.91 ± 0.22 kPa, 7.02 ± 0.31 kPa, and 7.64 ± 1.14 kPa for soft, intermediate, and stiff viscoelastic model ECMs, respectively. The measured $G_\infty$ values were 0.859 ± 0.12 kPa, 6.66 ± 0.186 kPa, and 7.37 ± 0.634 kPa for soft, intermediate, and stiff viscoelastic model ECMs, respectively. Results reveal that within the investigated times 1.03 – 1000 s, viscoelastic model ECMs did display evident relaxation behavior.

Figure S2c shows the average amplitude difference ($G_0$-$G_\infty$) for soft, intermediate, and stiff viscoelastic model ECMs. The average differences were ($G_0$-$G_\infty$) = 0.78 ± 0.19 kPa, ($G_0$-$G_\infty$) = 6.27 ± 0.31 kPa, ($G_0$-$G_\infty$) = 6.27 ± 0.71 kPa for soft, intermediate, and stiff viscoelastic model ECMs, respectively. It is interesting to note that the amplitude difference of soft is almost 8 times smaller than intermediate and stiff. Figure S2d shows the average relaxation times. These were $\tau$ = 1.47 ± 0.45 s, 2.90 ± 0.34 s, and 0.56 ± 0.11 s for soft, intermediate, and stiff, respectively.

**Table S1.**Values of relaxation modulus, G(t), obtained by relaxation test of elastic polyacrylamide model ECMs[40]

| Elastic | G ($t$=0.103) (kPa) | $G_\infty$ (kPa) | Amplitude decay, $G_\infty$-$G_0$ (kPa) | Relaxation time, τ (sec, s) |
|---|---|---|---|---|
| Soft E | 3.12 ± 0.00 | 3.14 ± 0.00 | -- | -- |
| Intermediate E | 8.15 ± 0.00 | 8.07 ± 0.00 | -- | -- |
| Stiff E | 12.06 ± 0.00 | 11.94 ± 0.01 | -- | -- |

**Table S2.** Values of relaxation modulus, G(t), obtained by relaxation test of viscoelastic polyacrylamide model ECMs.

| Viscoelastic | G (t=1.03) (kPa) | $G_\infty$ (kPa) | Amplitude decay, $G_\infty$-$G_0$ (kPa) | Relaxation time, τ (sec, s) |
|---|---|---|---|---|
| Soft VE | 0.905 ± 0.000 | 0.85 ± 0.12 | 0.784± 0.109 | 1.469 ± 0.259 |
| Intermediate VE | 7.02 ± 0.180 | 6.66 ± 0.187 | 6.26 ± 0.178 | 2.89 ± 0.199 |
| Stiff VE | 8.13 ± 0.00 | 7.37± 0.634 | 6.27 ± 0.407 | 0.452 ± 0.064 |

#### 4. Mean Square Displacement (MSD) of A549s cells crawling on elastic and viscoelastic model ECMs

Here we provide a summary of our observations of the mean square displacement curves for A549s cells migrating over a period of 24 hours on elastic and viscoelastic model ECMs.

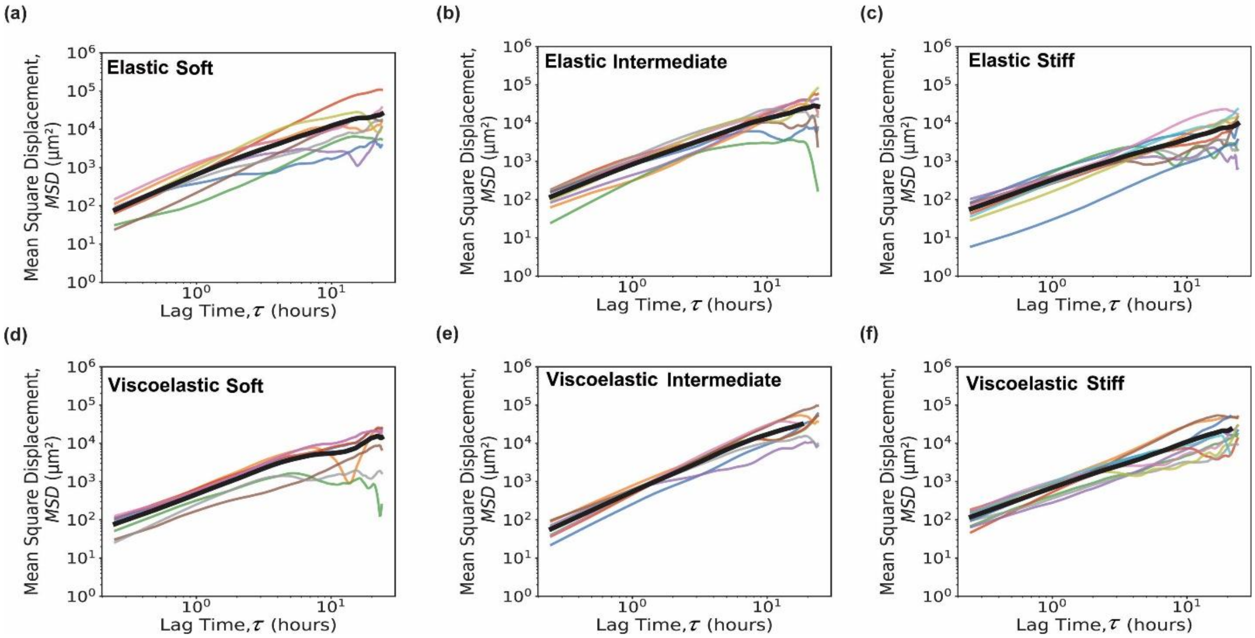


**Figure S4.** Total mean square displacement (MSD) versus lag time, τ, for A549 cells migrating on soft, intermediate, and stiff elastic and viscoelastic model ECMs over a period of 24 hours. Measurements were taken at 15-minute intervals. The mean square displacement was evaluated by tracking the centroid of the cells.

### 5. Average cell migration increments of 0-6, 6-12, 12-18, and 18-24 hours for A549 cells

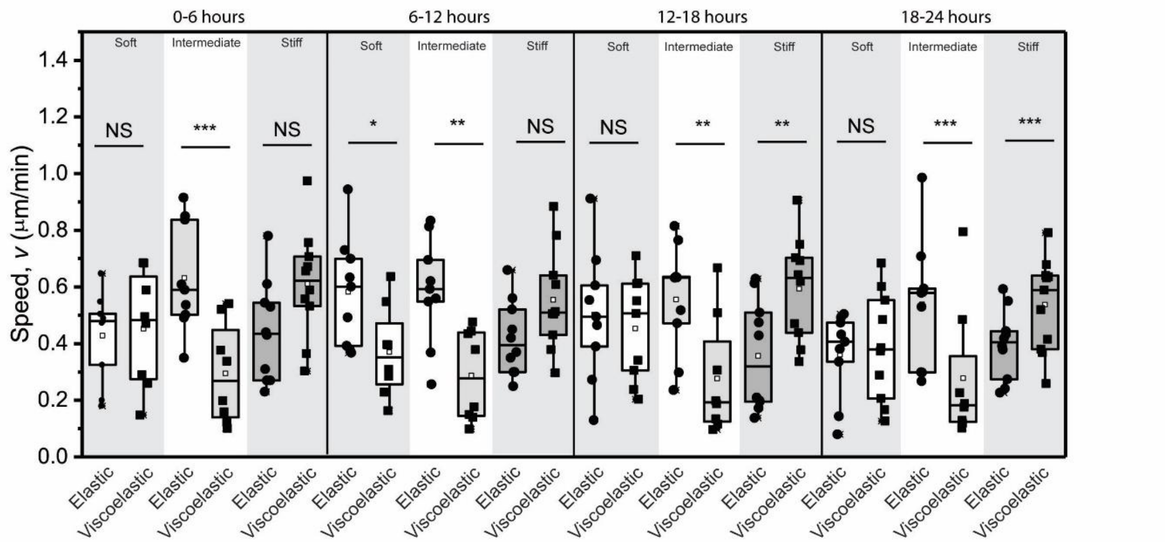


**Figure S5**. The migration speed of A549 cells on soft, intermediate, and stiff elastic and viscoelastic model ECMs over time increments of 0-6, 6-12, 12-18 and 18-24 hours, respectively. An independent t-test was used to determine if the differences in cell behavior between elastic and viscoelastic model ECMs were statistically significant. Notation: NS - not significant, * $p < 0.05$, ** $p < 0.01$, *** $p < 0.001$.

Cell speed was measured over the 24-hour period and analyzed across three intervals: 0-6 hours, 6-12 hours, and 12-24 hours. These results were compared with the overall results evaluated for the complete 0-24-hour period to assess whether migration behavior remains similar. Figure S4 shows the average speed of A549 on collagen type-I in different time periods, 0-6 hours, 6-12, and 12-24 hours on soft, intermediate, and stiff elastic and viscoelastic ECM models. Overall, cells on elastic ECMs exhibited similar migration rates over 24 hours. The average speeds over 0 - 6 hours was evaluated to be 0.42 ± 0.053 **μ**m/min, 0.63 ± 0.06 **μ**m/min, and 0.44 ± 0.05 **μ**m/min for elastic soft, intermediate, and stiff ECMs respectively, and 0.45 ± 0.07 **μ**m/min, 0.29 ± 0.06 **μ**m/min, and 0.61 ± 0.064 **μ**m/min for viscoelastic soft, intermediate, and stiff ECMs, respectively. The average speeds over the 6 to 12 hour period were 0.582 ± 0.069 **μ**m/min, 0.587 ± 0.067 **μ**m/min, 0.418 ± 0.04 **μ**m/min for the elastic ECMs, and 0.370 ± 0.056 **μ**m/min, 0.28 ± 0.05 **μ**m/min, 0.554 ± 0.06 **μ**m/min for the viscoelastic ESMs. The average speeds over the 12 -18 hours period were 0.502 ± 0.07 **μ**m/min, 0.55 ± 0.07 **μ**m/min, 0.357 ± 0.06 **μ**m/min for elastic ECMs and 0.45 ± 0.06 **μ**m/min, 0.33 ± 0.09 **μ**m/min, 0.59 ± 0.06 **μ**m/min for viscoelastic ECMs in comparison to 18-24: 0.36 ± 0.01 **μ**m/min, 0.53 ± 0.02 **μ**m/min, 0.39 ± 0.01 **μ**m/min Elastic and 0.38 ± 0.01 **μ**m/min, 0.32 ± 0.03 **μ**m/min, and 0.53 ± 0.01 **μ**m/min viscoelastic.

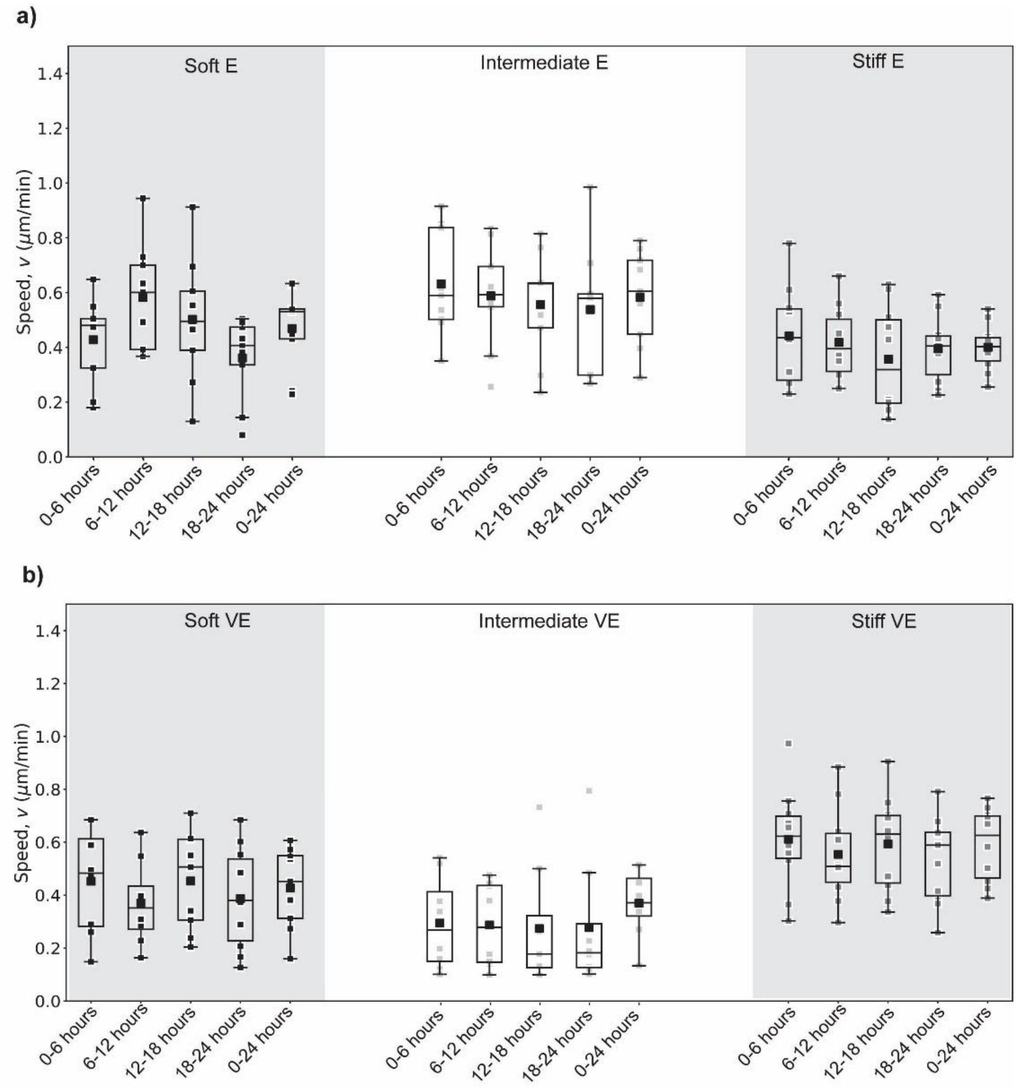


**Figure S6**. (a) A549 cell migration speed was grouped into classes based on time increments over 24 hours on soft, intermediate, and stiff elastic model ECMs, respectively. (b) A549 cell migration speed was grouped in different time increments over 24 hours on soft, intermediate, and stiff viscoelastic model ECMs, respectively.

### 6. UV-Vis of viscoelastic model ECMs

To determine the optical properties of viscoelastic polyacrylamide model ECMs, UV-Vis spectroscopy (measured using Aquamate 8000, ThermoScientific) of fully cured PAHs was performed. The absorbance, *A*, was measured for wavelengths between 200 and 1100 nm.

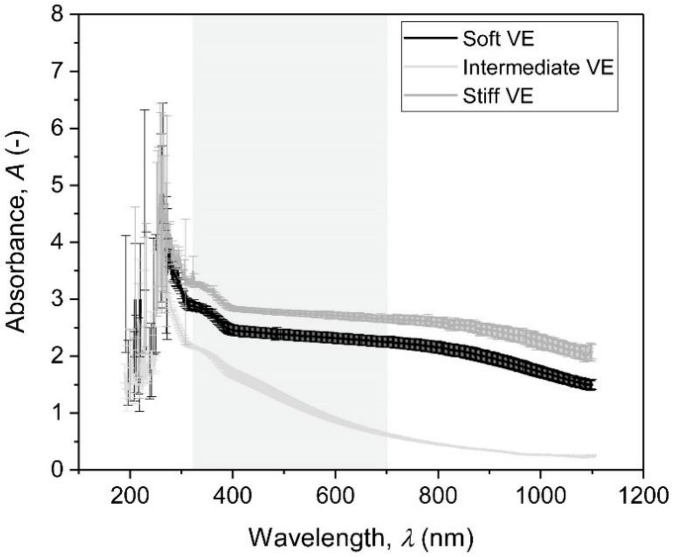


**Figure S7**. UV-Vis spectra comparison of Polyacrylamide soft, intermediate, and stiff viscoelastic model ECMs.

### 7. Additional cell migration speed over several time increments

Cell speed was observed throughout the 24 hours in different time increments, 0-6 hours, 6-12 hours, and 12-24 hours in comparison to 0-24 hours to see if migration behavior remains similar. Figure S6 a and b show the average speed of A549 on collagen type-I in different time increments, 0-6 hours, 6-12, and 12-24 hours on soft, intermediate, and stiff elastic and viscoelastic ECM models. Overall, cells on elastic ECM showed a similar migration speed over 24 hours. The measured speed from 0 - 6 hours was 0.42 ± 0.05 ***µ***m/min, 0.63 ± 0.06 ***µ***m/min, and 0.44 ± 0.05 ***µ***m/min for elastic and 0.458 ± 0.07 ***µ***m/min, 0.29 ± 0.06 ***µ***m/min, and 0.61 ± 0.06 ***µ***m/min for viscoelastic soft, intermediate, and stiff ECMs, respectively. The measured speed from 6.25-12 hours: 0.582 ± 0.06 ***µ***m/min, 0.587 ± 0.06 ***µ***m/min, 0.418 ± 0.041 ***µ***m/min elastic, and 0.370 ± 0.056 ***µ***m/min, 0.287 ± 0.05 ***µ***m/min, 0.55 ± 0.056 ***µ***m/min viscoelastic. The measured speed from 12-18 hours: 0.50 ± 0.07 ***µ***m/min, 0.55 ± 0.065 ***µ***m/min, 0.357 ± 0.06 ***µ***m/min elastic and 0.453 ± 0.06 ***µ***m/min, 0.27 ± 0.079 ***µ***m/min, 0.594 ± 0.058 ***µ***m/min viscoelastic The measured speed from 18-24 hours: 0.36 ± 0.05 ***µ***m/min, 0.538 ± 0.07 ***µ***m/min, 0.395 ± 0.03 ***µ***m/min elastic and 0.38 ± 0.06 ***µ***m/min, 0.27 ± 0.085 ***µ***m/min, 0.537 ± 0.049 ***µ***m/min viscoelastic in comparison to 0-24: 0.47 ± 0.05 ***µ***m/min, 0.58 ± 0.06 ***µ***m/min, 0.40 ± 0.027 ***µ***m/min Elastic and 0.43 ± 0.05 ***µ***m/min, 0.37 ± 0.04 ***µ***m/min, and 0.58 ± 0.04 ***µ***m/min viscoelastic.
